# Engineered vasculature induces functional maturation of pluripotent stem cell-derived islet organoids

**DOI:** 10.1101/2022.10.28.513298

**Authors:** Kim-Vy Nguyen-Ngoc, Yesl Jun, Somesh Sai, R. Hugh F. Bender, Vira Kravets, Han Zhu, Christopher J. Hatch, Michael Schlichting, Roberto Gaetani, Medhavi Mallick, Stephanie J. Hachey, Karen L. Christman, Steven C. George, Christopher C.W. Hughes, Maike Sander

## Abstract

Blood vessels play a critical role in pancreatic islet health and function, yet current culture methods to generate islet organoids from human pluripotent stem cells (SC-islets) lack a vascular component. Here, we engineered 3D vascularized SC-islet organoids by assembling SC-islet cells, human primary endothelial cells (ECs) and fibroblasts both in a non-perfused model and a microfluidic device with perfused vessels. Vasculature improved stimulus-dependent Ca^2+^ influx into SC-β-cells, a hallmark of β-cell function that is blunted in non-vascularized SC-islets. We show that an islet-like basement membrane is formed by vasculature and contributes to the functional improvement of SC-β-cells. Furthermore, cell-cell communication networks based on scRNA-seq data predicted BMP2/4-BMPR2 signaling from ECs to SC-β-cells. Correspondingly, BMP4 augmented the SC-β-cell Ca^2+^ response and insulin secretion. These vascularized SC-islet models will enable further studies of crosstalk between β-cells and ECs and can serve as *in vivo*-mimicking platforms for disease modeling and therapeutic testing.

## Introduction

Organoid models have become increasingly important for studying human organ development and modeling human diseases [1], as it has become clear that 3D organoids better mimic cellular functions in native tissues than traditional 2D culture models [2]. In the organ niche, supporting cells, such as stromal and vascular endothelial cells (ECs), provide important paracrine signals for cell maturation and function [3, 4]. Therefore, to create a model that fully mimics the *in vivo* environment, it is important to generate organoids that recapitulate the cellular complexity and physiological microenvironment of the organ niche [5, 6]. Furthermore, cells experience fluid and blood flow in their native tissue environment, creating mechanical forces that impact cellular states. These forces can be created *ex vivo* by culturing organoids in microphysiological systems, also known as organ-on-a-chip technology. The fluid flow introduced by microphysiological systems has been shown to promote cell maturation in human pluripotent stem cell-derived organoids [7, 8].

Methodology has been developed to generate 3D clusters akin to pancreatic islets from human pluripotent stem cells [9-12]. These stem cell-derived islets (SC-islets) comprise insulin-producing β-cells, glucagon-producing α-cells, and somatostatin-producing δ-cells. However, despite recent improvements in differentiation protocols, *in vitro* stem cell-derived β-cells (SC-β-cells) are functionally immature and do not respond to insulin secretory signals in the same manner as primary β-cells [10]. SC-β-cells acquire a more mature state after *in vivo* engraftment [10, 13], implying that the *ex vivo* culture model lacks important environmental cues for β-cell maturation present *in vivo*. An SC-islet model that incorporates non-endocrine cell types that are part of the *in vivo* tissue environment is still lacking.

In native islets, hormone-producing endocrine cells are surrounded by a vascular network of ECs, stromal cells, neurons, and immune cells. The dense vascular network that surrounds and penetrates each islet enables proper β-cell glucose sensing and insulin secretion into the circulation [14]. By secreting extracellular matrix (ECM) components and soluble factors, ECs establish a local microenvironment in the islet that provides important signals for β-cell function [15, 16]. Direct contact of β-cells with basement membrane ECM in the vascular bed stimulates integrin-β1 signaling to regulate various aspects of β-cell function including insulin production [17], Ca^2+^ influx [18], and insulin exocytosis [19]. In addition, EC-derived soluble factors provide important paracrine signals for β-cell function [20, 21]. During the isolation of islets from the pancreas and subsequent *in vitro* culture, islet vasculature is rapidly lost, which is a major contributor to impaired β-cell function in cultured islets [22]. Despite the importance of vasculature for islet function, there is currently no *in vitro* model that incorporates a 3D vasculature network to replicate the microenvironment established by ECs in the native islet niche.

Here, we established a vascularized SC-islet organoid model in which ECs form a vascular network and establish a basement membrane-like structure surrounding β-cells. To monitor β-cell function in the organoid, we introduced a GCaMP6f reporter into human pluripotent stem cells and quantified Ca^2+^ signals as a surrogate for insulin secretion in SC-β-cells. We find that the presence of a vascular network improves β-cell functional maturity in SC-islets. Furthermore, by establishing the vascularized SC-islet organoid in a microphysiological system, we show that β-cell function is further improved in the presence of fluid flow through the vascular network. Finally, we probed cell-cell communication networks in the organoid based on single-cell RNA-seq analysis, identifying ECM-β-cell signaling and BMP4 signaling from ECs to β-cells as important signals for improved β-cell function. Our organoid model highlights the importance of vasculature for maturing SC-β-cells *in vitro* and paves the way for improved modeling of human islet function and disease.

## Results

### Establishment of co-culture conditions for SC-islets with vascular endothelial cells

Routinely, ECs and SC-islets each require a specific culture medium for optimal cell viability. To establish a co-culture method for SC-islets and ECs, we tested how each medium affects survival of the other cell type by exposing SC-islets or ECs to the medium for the other cell type or mixtures between the two media (**Fig. S1A,B** and **Table S1**). We assessed EC integrity by their ability to form a vascular network when embedded with fibroblasts (FBs) into a 3D fibrin gel. Culture of SC-islets in ≥75% vascular medium significantly reduced the percentage of SC-β-cells (**Fig. S1C,D**) as well as insulin content of SC-islets (**Fig. S1E**). Conversely, <75% of vascular medium negatively affected vascular network formation and caused gel contraction (**Fig. S1F,G**). Thus, 75% of vascular medium mixed with 25% of islet medium supports vascular network formation but diminishes insulin production by SC-β-cells.

We next tested whether a gradual decrease of vascular medium over time could maintain both SC-β-cell insulin production and a vascular network (**Fig. S1H**). A gradual switch from 75% vascular plus 25% islet medium to 100% islet medium - referred to as gradual medium change 1 (GM1) condition (**Table S1**) - was most effective in preserving insulin content of SC-islets (**Fig. S1I**). The GM1 condition also maintained a vascular network, albeit compromised when compared to the network in 100% vascular medium (**Fig. S1J,K**). Together, these experiments identify a mixed 75% vascular and 25% islet medium as most effective in preserving a vascular network and a gradual change into 100% islet medium as optimal for maintaining SC-β-cell insulin content.

To establish a co-culture model of SC-islets and ECs, we tested (i) 75% of vascular medium mixed with 25% of islet medium and (ii) GM1 gradual medium change into islet medium in co-cultures of SC-islets, ECs, and FBs (**Fig. 1A**). To this end, we resized SC-islets to <200 μm, embedded SC-islets with ECs and FBs into 3D fibrin gels and cultured the aggregates for five days (**Fig. 1A**). Whereas ECs and FBs gradually exposed to 100% islet medium poorly formed vascular networks (**Fig. 1C** and **S1J,K**), ECs and FBs co-cultured with SC-islets maintained a fully developed vascular network after gradual change into islet medium (**Fig. 1B-D**). Consistent with findings for SC-islets cultured alone (**Fig. S1I**), this condition allowed for maintenance of insulin content also in vascularized SC-islets (**Fig. 1E**). In vascularized SC-islet organoids, vasculature tightly wrapped around SC-islets and ECs were in the immediate vicinity of SC-β-cells; however, the vasculature rarely penetrated through SC-islets (**Fig. 1F**). These results establish a culture condition for vascularizing SC-islets.

**Figure 1.**
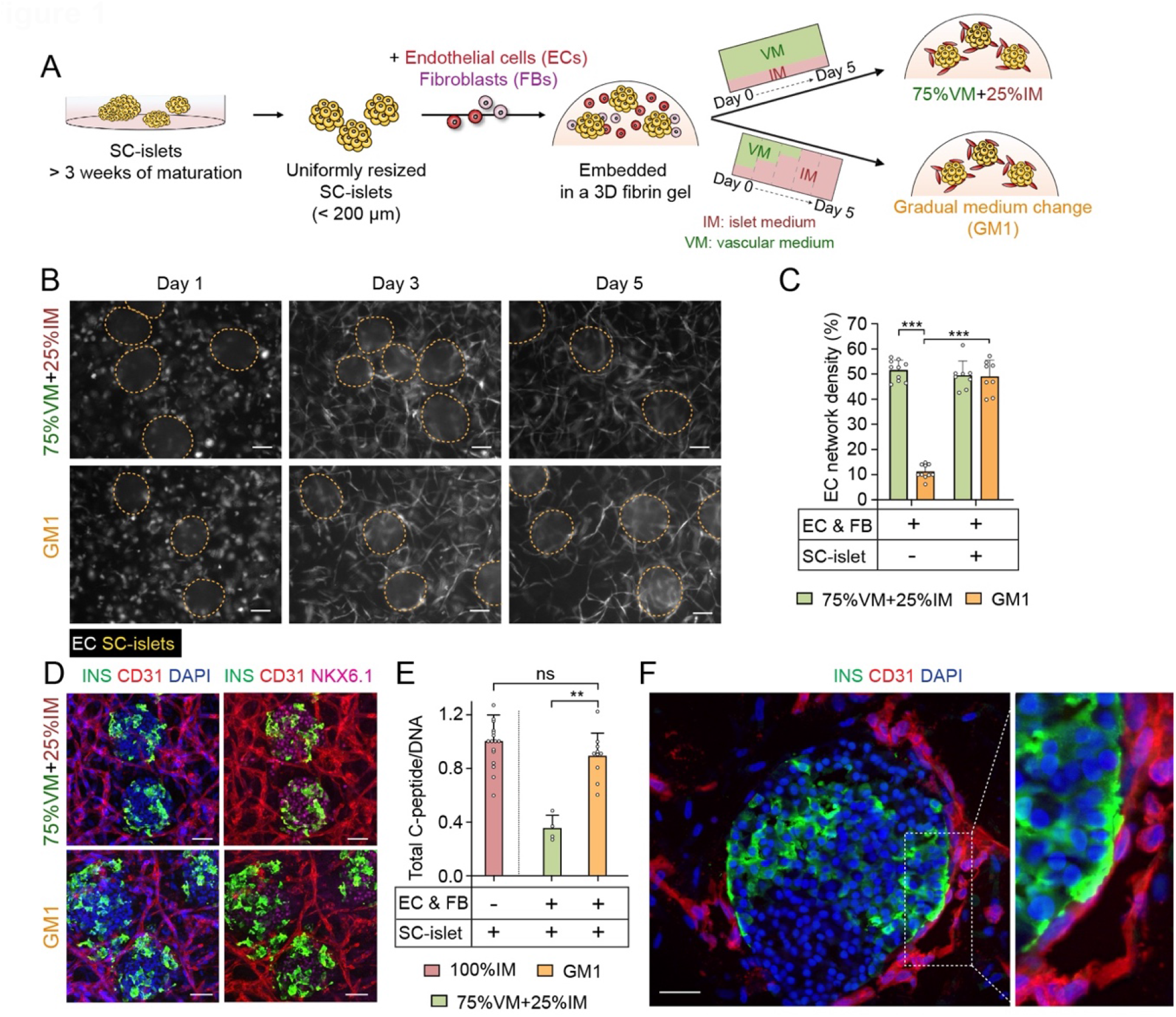
Establishment of a vascularized SC-islet organoid. (**A**) Schematic depicting establishment of the method for vascularizing SC-islets. (**B**) Representative epifluorescence images of SC-islets and endothelial cells (EC) co-cultured in 75%VM + 25%IM or GM1. ECs were transduced with an azurite blue-expressing lentivirus. SC-islets are circled. Scale bars, 50 μm. (**C**) Quantification of vascular network density at day 5 of co-culture in 75%VM + 25%IM or GM1 (*n* = 10 for culture without SC-islets, and *n* = 8 for culture with SC-islets from 3 independent experiments). Dots denote data points of individual 3D fibrin gels. (**D**) Representative immunofluorescence images of vascularized SC-islets stained with insulin (INS), CD31, and NKX6.1 after culture in 75%VM + 25%IM or GM1 for 5 days. Scale bars, 50 μm. (**E**) Quantification of C-peptide content in vascularized SC-islets cultured in 75%VM + 25%IM or GM1, and non-vascularized SC-islets cultured in 100% IM for 5 days (*n* = 17 for IM, *n* = 10 for GM1, and *n* = 4 for 75%VM + 25%IM from 10, 6, and 3 independent experiments, respectively). Dots denote data points of individual 3D fibrin gels containing 50 SC-islets each. (**F**) Representative immunofluorescence image of vascularized SC-islets stained with INS and CD31. The inset shows close contact of SC-β-cells with ECs. Scale bar, 30 μm. All data are shown as mean ± S.D., Student’s *t* test, **p-value < 0.01, ***p-value < 0.001, ns, not significant.

### A GCaMP6f-hESC reporter line for calcium signal recording in SC-β-cells

In native islets, ECs play an important role in the control of insulin secretion by β-cells [15]. In the vascularized SC-islet organoid model, insulin secretion cannot be reliably measured because the fibrin gel constitutes a barrier for insulin release into the culture medium. Therefore, we generated a human embryonic stem cell line (hESCs) carrying the genetically encoded green fluorescence calcium indicator GCaMP6f for monitoring insulin secretion. hESC-GCaMP6f differentiated into SC-islets comprising all major endocrine cell types in similar proportions as in SC-islets from the parent hESC line (**Fig. S2A-C**). Furthermore, SC-islets derived from hESC-GCaMP6f and the parent hESC line exhibited similar insulin secretory responses to glucose and the GLP-1 analogue Exendin-4 (**Fig. S2D)**, which augments glucose-mediated insulin secretion by promoting calcium (Ca^2+^) influx [23]. Thus, insertion of the GCaMP6f reporter did not change the properties of the hESC line.

To study the Ca^2+^ response of β-cells after stimulation with high glucose alone or Exendin-4 in the presence of high glucose, we established an imaging workflow for quantifying the intracellular Ca^2+^ signal of SC-β-cells in GCaMP6f-SC-islets. Previous work has shown that SC-β-cells can be identified by live staining for the cell surface marker CD49a [11], which we validated in our system (**Fig. 2A**). To analyze Ca^2+^ signals in SC-β-cells, we combined Ca^2+^ signal recording with staining for CD49a, insulin (INS) and NKX6.1 (**Fig. 2A**). In total, we analyzed Ca^2+^ traces of 211 SC-β-cells from seven different SC-islets exposed to low glucose, high glucose, Exendin-4, and KCl, which triggers membrane depolarization and Ca^2+^ influx (**Fig. 2B,C** and **Video S1**). Based on their Ca^2+^ response to glucose and Exendin-4, we classified SC-β-cells into four groups: (i) responsive to both high glucose and Exendin-4, (ii) responsive to only high glucose but not Exendin-4, (iii) responsive to Exendin-4 but not high glucose, and (iv) non-responsive (**Fig. 2D** and **S2E**). We found that only 17 ± 12.5% of SC-β-cells exhibited a Ca^2+^ response to both high glucose and Exendin-4, while 51.5 ± 25% of SC-β-cells were non-responsive (**Fig. 2D,E**). The amplitude of the SC-β-cell Ca^2+^ response to high glucose was small (**Fig. 2D** and **S2E**), consistent with a prior study that compared Ca^2+^ signals in SC-β-cells and primary islet β-cells [10].

**Figure 2.**
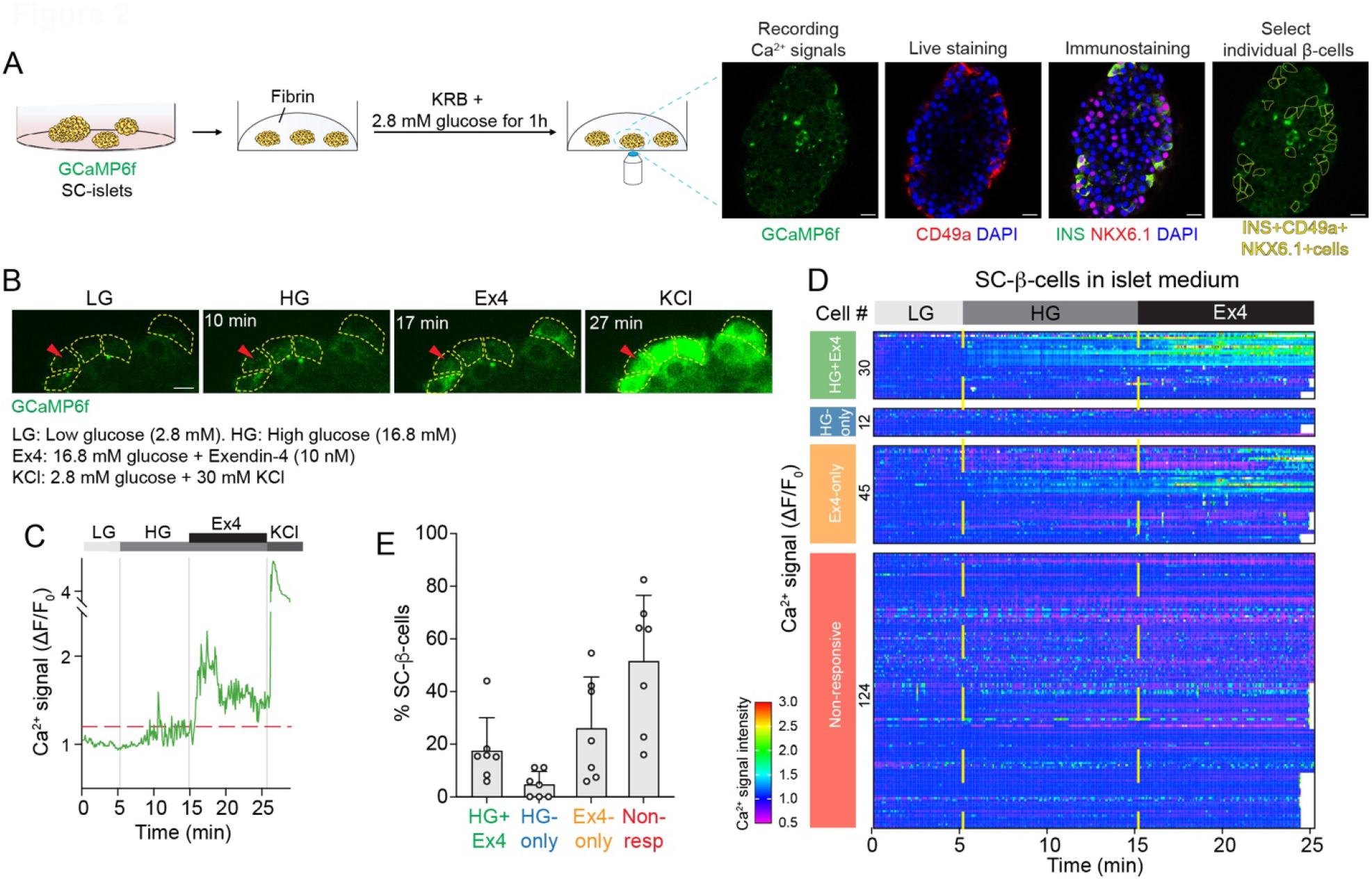
Stimulus-dependent Ca^2+^ responses in GCaMP6f-SC-β-cells. (**A**) Schematic depicting workflow to measure intracellular Ca^2+^ signals in GCaMP6f**-**SC-β-cells. Scale bars, 50 μm; insulin, INS. (**B**) Still images of Ca^2+^ signals of select SC-β-cells sequentially exposed to 2.8 mM glucose (LG), 16.8 mM glucose (HG), 10 nM Exendin-4 (Ex4), and 30 mM KCl. Scale bar, 10 μm. Arrowhead points to an SC-β-cell, of which Ca^2+^ trace is shown in (C). (**C**) Representative trace of Ca^2+^ signal in one SC-β-cell shown in (B) in response to depicted stimuli. (**D**) Heatmap showing normalized Ca^2+^ signals of 211 SC-β-cells from 7 SC-islets exposed to LG, HG, and Ex4. High Ca^2+^ signal (red), low Ca^2+^ signal (violet). (**E**) Quantification of SC-β-cells exhibiting a Ca^2+^ response to HG and Ex4, HG but not Ex4 (HG-only), Ex4 but not HG (Ex4-only) or being non-responsive to either HG or Ex4 (Non-resp) (*n* = 7 SC-islets from 3 independent experiments; dots denote data points of individual SC-islets; data are shown as mean ± S.D.).

### SC-islet vascularization augments the SC-β-cell Ca^2+^ response

Having established Ca^2+^ signal recording in SC-β-cells, we next sought to compare the Ca^2+^ response of SC-β-cells in the non-vascularized and vascularized SC-islet models. To this end, we cultured GCaMP6f-SC-islets with or without ECs and FBs, switching gradually from 75% vascular plus 25% islet medium at day 1 to 100% islet medium before recording Ca^2+^ traces in SC-β-cells at day 5 (**Fig. 3A**). Comparative analysis of SC-β-cells from non-vascularized and vascularized SC-islets revealed a significant increase in the percentage of SC-β-cells exhibiting a Ca^2+^ response to both high glucose and Exendin-4 in the vascularized SC-islet model (**Fig. 3B-D, S3A,B**, and **Videos S2** and **S3**). Correspondingly, the percentage of non-responsive SC-β-cells was lower in the vascularized model. Furthermore, quantification of Ca^2+^ traces through area under the curve (AUC) measurement showed that the magnitude of the SC-β-cell Ca^2+^ response after glucose and Exendin-4 stimulation was greater in vascularized than in non-vascularized SC-islets (**Fig. 3E**), while the peak height of the Ca^2+^ signal did not differ in the two models (**Fig. 3F**). Together, these findings demonstrate that co-culture of SC-islets with microvessels enhances the Ca^2+^ response of SC-β-cells upon glucose and Exendin-4 stimulation, suggesting that inclusion of vessels into SC-islet organoids provides important functional cues for SC-β-cells.

**Figure 3.**
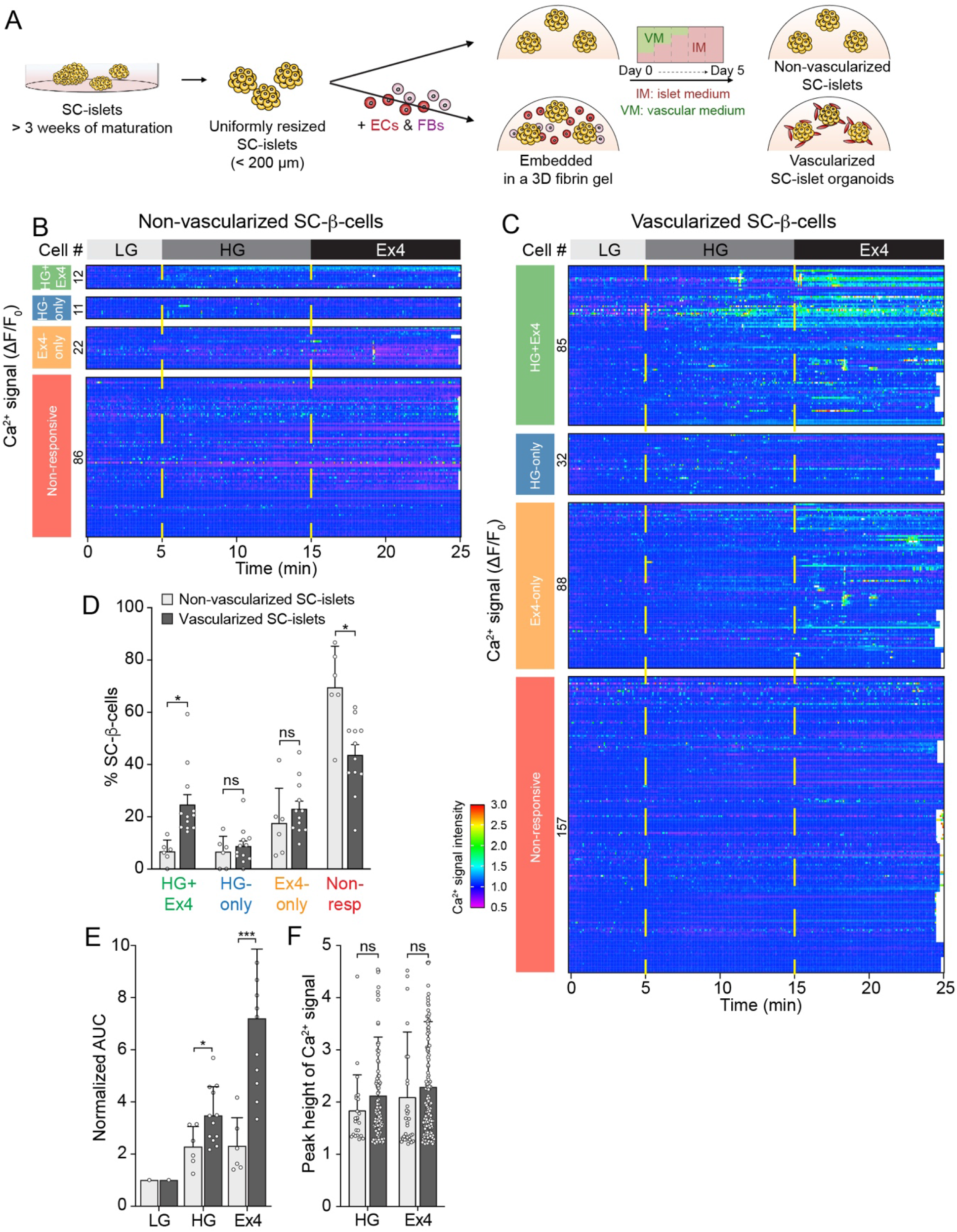
Improved Ca^2+^ response of SC-β-cells after SC-islet vascularization. (**A**) Schematic depicting generation of non-vascularized and vascularized SC-islet organoids. (**B** and **C**) Heatmaps showing normalized Ca^2+^ signals of 131 SC-β-cells from 6 non-vascularized SC-islets (B) and 362 SC-β-cells from 12 vascularized SC-islets (C) exposed to LG, HG, and Ex4. High Ca^2+^ signal (red), low Ca^2+^ signal (violet). (**D**) Quantification of SC-β-cells exhibiting a Ca^2+^ response to HG and Ex4, HG but not Ex4 (HG-only), Ex4 but not HG (Ex4-only) or being non-responsive to either HG or Ex4 (Non-resp) in non-vascularized and vascularized SC-islets (*n* = 6 non-vascularized and *n* = 12 vascularized SC-islets from 4 and 7 independent experiments, respectively; dots denote data points of individual SC-islets). (**E**) Quantification of area under the curve (AUC) of SC-β-cells exhibiting a Ca^2+^ response in HG (from HG+Ex4 and HG-only groups) or Ex4 (from HG+Ex4 and Ex4-only groups). *n* = 6 non-vascularized and *n* = 12 vascularized SC-islets from 4 and 7 independent experiments, respectively; dots denote data points of individual SC-islets. The AUC value in LG was set as 1. (**F**) Quantification of Ca^2+^ signal peak height of responsive SC-β-cells in HG (from HG+Ex4 and HG-only groups; *n* = 23 SC-β-cells in non-vascularized and *n* = 117 SC-β-cells in vascularized SC-islets) or Ex4 (from HG+Ex4 and Ex4-only groups; *n* = 34 SC-β-cells in non-vascularized and *n* = 173 SC-β-cells in vascularized SC-islets). Dots denote data points of individual SC-β-cells. All data are shown as mean ± S.D., Student’s *t* test, *p-value < 0.05, ***p-value < 0.001, ns, not significant.

### A perfusable vascular network prolongs Ca^2+^ influx into SC-β-cells

Under physiological conditions, the endothelium is exposed to continuous shear stress due to interstitial and pulsatile blood flow, which has been shown to stimulate vascular network formation [24] and islet function [25]. To introduce vessel flow into vasculature surrounding SC-islets, we adapted a vascularized micro-organ microfluidic system, in which ECs and FBs are seeded to form a network of perfused microvessels in a microfluidic device [25-27]. In this system, the shear of interstitial flow induces formation of a vascular network that is perfused through the microfluidic channels (**Fig. S4A**). Using the condition of gradual change from vascular into islet medium (**Table S2**), we loaded SC-islets together with ECs and FBs into the microfluidic device (**Fig. 4A**). After six days, a perfusable vascular network surrounding SC-islets formed inside the tissue chamber (**Fig. 4B,C** and **S4B**). SC-islets inside the device maintained robust expression of insulin (**Fig. 4C**). Quantification of Ca^2+^ responsive SC-β-cells in the device (**Fig. S4C-E** and **Video S4**) identified 26 ± 15.3% of SC-β-cells responding to both high glucose and Exendin-4 (**Fig. 4D,E** and **S4F**), which is a similar percentage as in the vascularized SC-islet model outside the microfluidic device (**Fig. 3C,D**). Quantification of Ca^2+^ traces revealed an AUC of 7.4 ± 3.1 in high glucose and 25.7 ± 13.5 in Exendin-4 in the microfluidic device (**Fig. 4F**) compared to 3.5 ± 1.1 and 7.2 ± 2.7, respectively, outside the device (**Fig. 3E**). This analysis suggests prolonged post-stimulation Ca^2+^ influx in vascularized SC-β-cells in the device compared to regular cultures. Peak height of the Ca^2+^ signal did not differ between the regular culture model (**Fig. 3F**) and the device (**Fig. 4G**). The results demonstrate successful incorporation of SC-islets into a perfusable vasculature in a microfluidic device and suggest that the perfusable system augments the duration of Ca^2+^ influx into SC-β-cells after stimulation.

**Figure 4.**
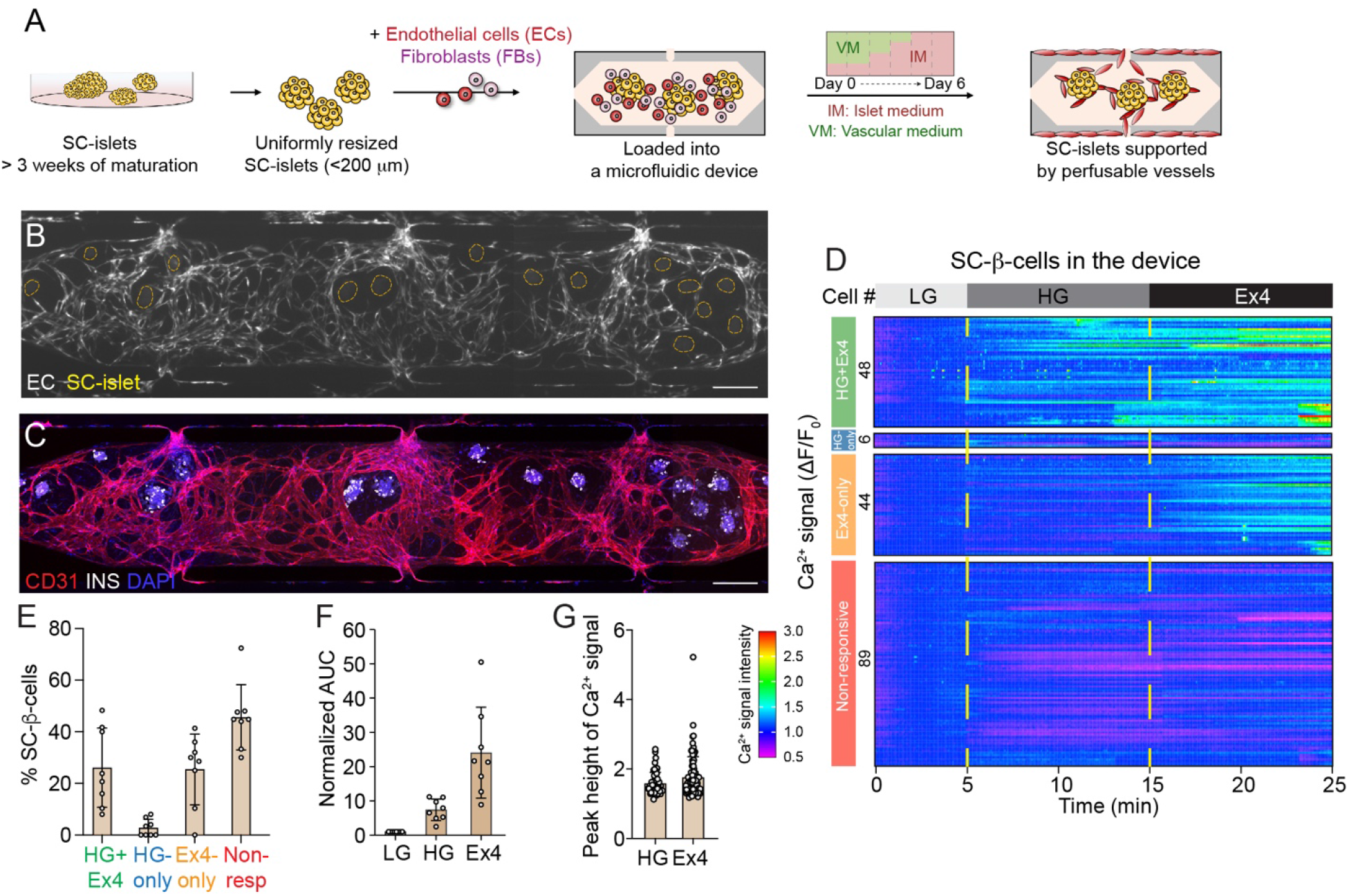
Vascularized SC-islet organoids in a microfluidic device. (**A**) Schematic depicting incorporation of vascularized SC-islet organoids into microfluidic device. (**B**) Representative epifluorescence image of live vascular network on day 6 in the device. ECs were transduced with an azurite blue-expressing lentivirus. SC-islets are circled. Scale bar, 200 μm. (**C**) Representative immunofluorescence image of vascularized SC-islet organoids on day 6 in the device stained with insulin (INS), CD31, and DAPI. Scale bar, 200 μm. (**D**) Heatmap showing normalized Ca^2+^ signals of 187 SC-β-cells from 8 vascularized SC-islets exposed to LG, HG, and Ex4 in the device. High Ca^2+^ signal (red), low Ca^2+^ signal (violet). (**E**) Quantification of SC-β-cells exhibiting a Ca^2+^ response to HG and Ex4, HG but not Ex4 (HG-only), Ex4 but not HG (Ex4-only) or being non-responsive to either HG or Ex4 (Non-resp) in the device (*n* = 8 SC-islets from 6 devices; dots denote data points of individual SC-islets). (**F**) Quantification of area under the curve (AUC) of SC-β-cells exhibiting a Ca^2+^ response in HG (from HG+Ex4 and HG-only groups) or Ex4 (from HG+Ex4 and Ex4-only groups). *n* = 8 SC-islets from 6 devices; dots denote data points of individual SC-islets. The AUC value in LG was set as 1. (**G**) Quantification of Ca^2+^ signal peak height of responsive SC-β-cells in HG (from HG+Ex4 and HG-only groups; *n* = 54 SC-β-cells) or Ex4 (from HG+Ex4 and Ex4-only groups; *n* = 92 SC-β-cells). Dots denote data points of individual SC-β-cells. All data are shown as mean ± S.D.

### Prediction of cell-cell signaling networks in vascularized SC-islets

Our results suggest that the presence of a vascular network provides molecular cues that promote Ca^2+^ influx in SC-β-cells under stimulatory conditions. To identify signals that mediate these effects, we performed single-cell RNA-sequencing (scRNA-seq) of SC-islets and microvessels formed from ECs and FBs in the microfluidic device (**Table S3a**). From these data, we sought to identify candidate ligand-receptor pairs mediating communication between SC-β-cells, ECs and FBs. After applying quality control filters, 15,941 SC-islet cells and 1,809 vasculature-forming cells (ECs and FBs) were retained and subsequently integrated into a single Uniform Manifold Approximation and Projection (UMAP) (**Fig. 5A,B**). Cell type annotation based on known marker genes revealed distinct sub-clusters for each endocrine cell type within the SC-islet cluster, including SC-β-cells (*INS, NKX6*.*1*), SC-α-cells (*GCG*) and SC-δ-cells (*SST*), whereas vasculature-forming cells clustered separately into ECs (*PECAM1*) and FBs (*COL6A1*) (**Fig. 5B** and **S5A**).

**Figure 5.**
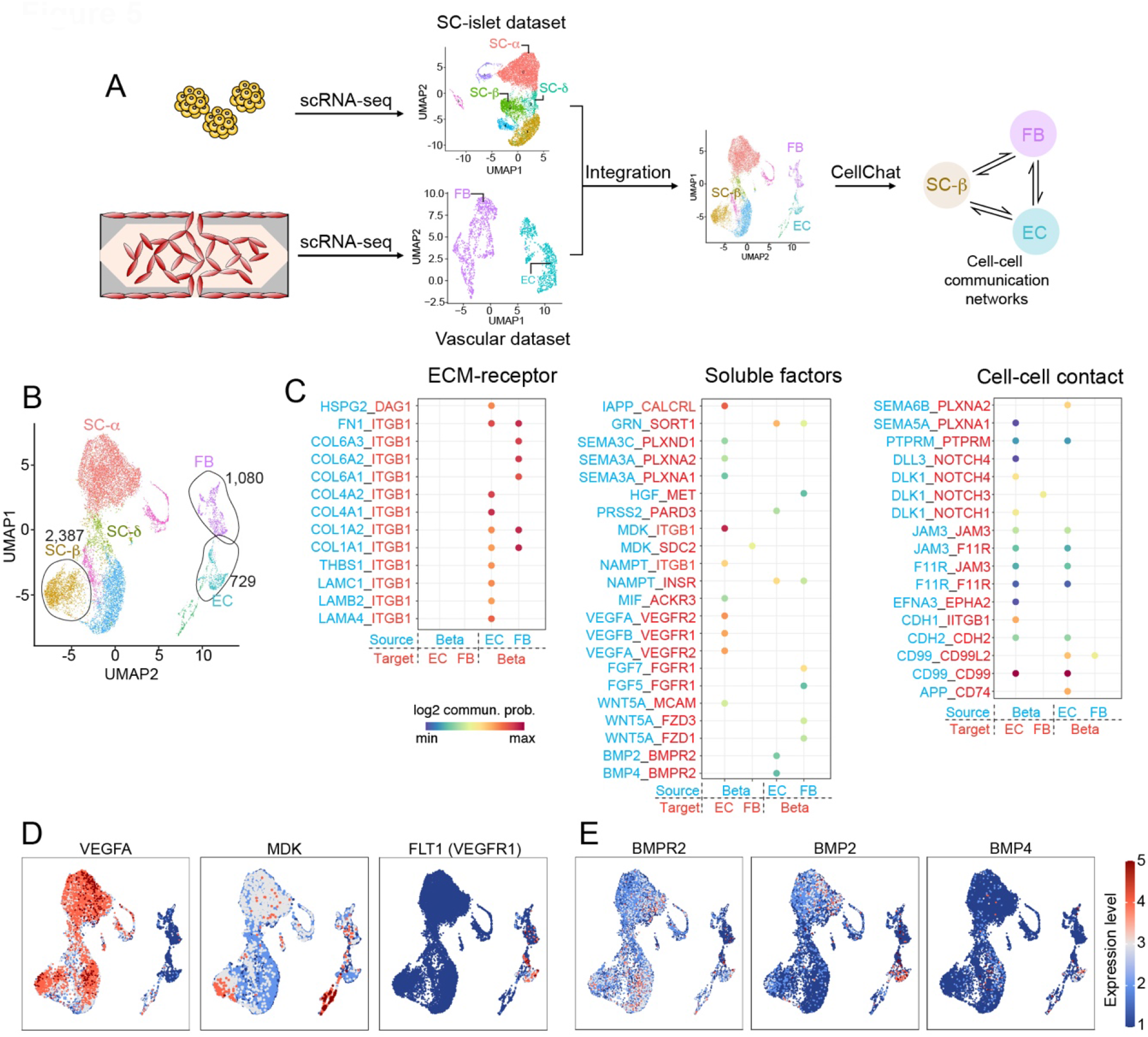
Predicted cell-cell communication networks between SC-β-cells and vasculature-forming cells. (**A**) Schematic of workflow for predicting cell-cell communication networks based on scRNA-seq data. FB, fibroblast; EC, endothelial cell; SC, stem cell. (**B**) Clustering of 15,941 SC-islet cells and 1,809 vasculature-forming cells. (**C**) Significant ligand-receptor pairs between SC-β-cells and ECs or FBs. Dot color reflects communication probabilities. (**D** and **E**) Feature plots showing expression levels of ligands and receptors for potential signals from SC-β-cells to ECs (D) and ECs to SC-β-cells (E). High expression (red) and low expression (blue).

To identify potential interactions between SC-β-cells, ECs and FBs, we generated a cell-cell communication network among SC-β-cells, ECs and FBs using CellChat [28], identifying a total of 333 possible interactions between the cell types (**Fig. S5B,C** and **Table S3b**). Most of the interactions were related to ECM-receptor signaling (**Fig. 5C** and **S5C**), consistent with the known role for islet vasculature in generating a local niche comprised of ECM proteins [17, 29]. FBs expressed collagen I, VI, and fibronectin and ECs expressed the basement membrane components collagen IV, laminin 1, 4, 5 (**Fig. S5A,D**), which are ligands for the integrin-β1 receptor (*ITGB1*) expressed by SC-β-cells (**Fig. S5D**). In addition, CellChat predicted 79 interactions via soluble factors between SC-β-cells and vasculature-forming cells (**Fig. 5C**). SC-β-cells expressed angiogenic factors *VEGFA* [30] and *MDK* [31], whereas ECs expressed the VEGF receptor (**Fig. 5D**), providing a possible mechanism for why co-culture of ECs with SC-islets improves vascular network density (**Fig. 1C**). FBs also expressed paracrine factors known to promote β-cell differentiation and maturation, including *WNT5A/WNT4* [32] and *HGF* [33], while SC-β-cells expressed the respective receptors (**Fig. S5E**). CellChat also predicted BMP2/4-BMPR2 signaling from ECs to SC-β-cells (**Fig. 5C,E**). Together, these data suggest that ECM to SC-β-cell signaling is a major communication pathway. In addition, we identify novel candidate signaling mechanisms (e.g., BMP signaling) between SC-β-cells and vasculature.

### Vasculature-derived ECM improves Ca^2+^ influx into SC-β-cells

In native islets, the basement membrane is predominantly comprised of ECM proteins produced by ECs [17]. To determine whether vascularization of SC-islets leads to formation of a structure akin to the islet basement membrane, we analyzed ECM protein expression in non-vascularized and vascularized SC-islets. Immunofluorescence staining for the basement membrane components laminin-α1 and collagen IV revealed a basement membrane-like structure surrounding vascularized SC-islets (**Fig. 6A**), similar to the peri-islet basement membrane [34, 35]. By contrast, non-vascularized SC-islets and cadaveric islets after the isolation procedure lacked this basement membrane-like structure (**Fig. S6A,B**). Consistent with a vascular origin of the ECM proteins, laminin-α1 and collagen IV mostly colocalized with CD31-positive ECs (**Fig. 6A**). Quantification of the percentage of laminin-α1- and collagen IV-positive cells in SC-islets showed more laminin-α1- and collagen IV-positive cells in vascularized compared to non-vascularized SC-islets and a higher proportion of SC-β-cells contacting ECM after vascularization (**Fig. 6A,B**). To determine whether SC-islet vascularization stimulates integrin signaling in SC-β-cells, we stained for activated integrin-β1. As predicted, vascularization increased the percentage of SC-β-cells with activated ECM-integrin signaling from 24.8 ± 9.4% to 49.1 ± 10.3% (**Fig. 6C,D**). To test whether activation of integrin signaling is mediated by basement membrane-like ECM, we encapsulated SC-islets into 3D gels comprised of fibrin alone or fibrin supplemented with Matrigel, which mimics basement membrane ECM components. Addition of Matrigel established a basement membrane-like structure surrounding SC-islets, activated integrin-β1 in SC-β-cells, and increased total C-peptide content of SC-islets (**Fig. 6E-G**).

**Figure 6.**
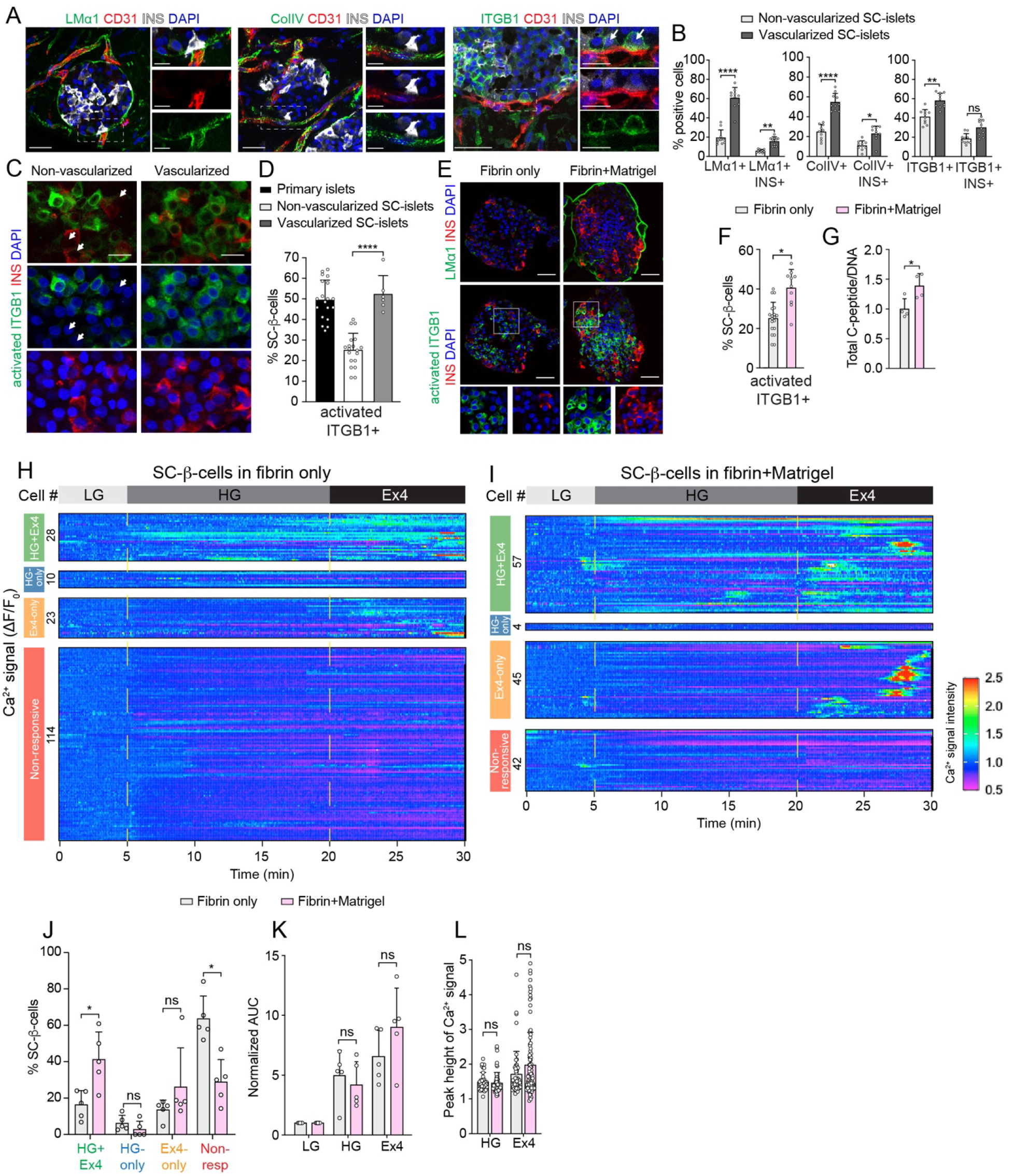
The ECM microenvironment of vascularized SC-islets improves SC-β-cell Ca^2+^ responsiveness. (**A**) Representative immunofluorescence images of peri-basement membrane-like structure in vascularized SC-islet organoids stained with laminin-α1 (LMα1), collagen IV (ColIV), integrin-β1 (ITGB1), CD31, and insulin (INS). Scale bars, 50 μm for large images and 20 μm for insets. Arrows point to INS^+^ cells expressing ITGB1. (**B**) Quantification of SC-β-cells in contact with basement membrane proteins (LMα1, ColIV) and expressing ITGB1 in non-vascularized and vascularized SC-islets (*n* = 10 from 3 independent experiments; dots denote data points of individual SC-islets). (**C**) Representative immunofluorescence images of non-vascularized and vascularized SC-islets stained with activated ITGB1, INS, and DAPI. Scale bars, 20 μm. Arrows point to INS^+^ cells lacking activated ITGB1 in non-vascularized SC-islets. (**D**) Quantification of SC-β-cells expressing activated ITGB1 in non-vascularized and vascularized SC-islets, and in primary human islets (*n* = 19 for both non-vascularized and vascularized SC-islets from 3 independent experiments, and *n* = 6 for primary human islets; dots denote data points of individual SC-islets and primary islets, respectively). (**E**) Representative immunofluorescence images of peri-basement membrane-like structure (top) and activated ITGB1 in SC-β-cells (bottom) of SC-islets encapsulated in 3D gels of fibrin only or fibrin+Matrigel (1:1) and stained for LMα1, activated ITGB1, and INS. Scale bars, 50 μm. Bottom images show boxed areas with split channels. (**F**) Quantification of SC-β-cells expressing activated ITGB1 in SC-islets encapsulated in 3D fibrin only or fibrin+Matrigel (1:1) (*n* = 19 for fibrin only and *n* = 10 for fibrin+Matrigel from 3 independent experiments; dots denote data points of individual SC-islets). (**G**) Quantification of C-peptide content in SC-islets encapsulated in 3D fibrin only or fibrin+Matrigel (1:1) (*n* = 4 gels from 3 independent experiments; dots denote data points of individual 3D gels containing 50 SC-islets each). (**H** and **I**) Heatmaps showing normalized Ca^2+^ signals of 175 SC-β-cells from 5 SC-islets encapsulated in 3D fibrin gels (H) and 148 SC-β-cells from 5 SC-islets encapsulated in fibrin+Matrigel (1:1) gels (I) exposed to LG, HG, and Ex4. High Ca^2+^ signal (red), low Ca^2+^ signal (violet). (**J**) Quantification of SC-β-cells exhibiting a Ca^2+^ response to HG and Ex4, HG but not Ex4 (HG-only), Ex4 but not HG (Ex4-only) or being non-responsive to either HG or Ex4 (Non-resp) in SC-islets encapsulated in 3D fibrin only or fibrin+Matrigel (1:1) (*n* = 5 for both fibrin only and fibrin+Matrigel SC-islets from 3 independent experiments; dots denote data points of individual SC-islets). (**K**) Quantification of area under the curve (AUC) of SC-β-cells exhibiting a Ca^2+^ response in HG (from HG+Ex4 and HG-only groups) or Ex4 (from HG+Ex4 and Ex4-only groups). *n* = 5 for both fibrin only and fibrin+Matrigel from 3 independent experiments; dots denote data points of individual SC-islets. The AUC value in LG was set as 1. (**L**) Quantification of Ca^2+^ signal peak height of responsive SC-β-cells in HG (from HG+Ex4 and HG-only groups; *n* = 38 SC-β-cells in fibrin only and *n* = 61 SC-β-cells in fibrin+Matrigel) or Ex4 (from HG + Ex4 and Ex-only groups; *n* = 51 SC-β-cells in fibrin only and *n* = 102 SC-β-cells in fibrin+Matrigel; dots denote data points of individual SC-β-cells). All data are shown as mean ± S.D., Student’s *t* test, *p-value < 0.05, **p-value < 0.01, ****p-value < 0.0001, ns, not significant.

Integrin signaling in β-cells enhances insulin secretion [17, 36]. To determine the effect of integrin-β1 activation on SC-β-cell function, we compared Ca^2+^ signals of SC-β-cells in SC-islets embedded in fibrin alone and in fibrin and Matrigel. Supplementation with Matrigel increased the percentage of SC-β-cells exhibiting a Ca^2+^ response to both high glucose and Exendin-4 but did not affect the magnitude of the SC-β-cell Ca^2+^ response or the average peak height of the signals (**Fig. 6H-L** and **Videos S5** and **S6**). This finding is consistent with enhanced glucose-dependent insulin secretion when SC-β-cells are cultured on basement membrane ECM proteins [37]. Together, these results show that basement membrane-like ECM produced by ECs in the vascularized SC-islet model augments integrin-β1 signaling in SC-β-cells and improves SC-β-cell function.

### BMP4 stimulates insulin secretion and Ca^2+^ influx in SC-β cells

The cell-cell signaling network also predicted BMP2-BMPR2 and BMP4-BMPR2 signaling from ECs to SC-β-cells (**Fig. 5C**). BMP2 and BMP4 are paralogs that both activate BMPR2 signaling [38, 39]. To determine how stimulation of BMPR2 signaling in SC-β-cells affects SC-β-cell function, we treated SC-islets with BMP4 across a wide range of dosages and measured glucose-stimulated insulin secretion and Ca^2+^ responses (**Fig. 7A**). As expected, BMP4 treatment activated canonical BMP signaling resulting in nuclear pSMAD1 accumulation in SC-β-cells (**Fig. S7A**). Measurement of glucose-stimulated insulin secretion in SC-islets revealed that intermediate BMP4 dosages (1, 5, or 10 ng/ml BMP4) significantly enhanced insulin secretion (**Fig. 7B**). Likewise, analysis of Ca^2+^ influx showed that treatment with 5 ng/ml BMP4 increased the percentage of SC-β-cells exhibiting a Ca^2+^ response to high glucose and Exendin-4 (**Fig. 7C-F, S7B-D**, and **Videos S7** and **S8**). In addition, BMP4 treatment robustly increased the peak height of the Ca^2+^ response (**Fig. 7G**). We further observed that Ca^2+^ oscillations appeared more synchronized between of SC-β-cells in BMP4-treated SC-islets (**Fig. 7D** and **S7B**). To quantify this, we conducted network analysis of Ca^2+^ dynamics [40] which revealed increased connectivity between BMP4-treated SC-β-cells at low and high glucose (**Fig. 7H**). These findings suggest that BMP4 plays a role in establishing β-β-cell functional coupling. Together, our results demonstrate the importance of paracrine signals from vascular ECs for the functional maturation of SC-β-cells.

**Figure 7.**
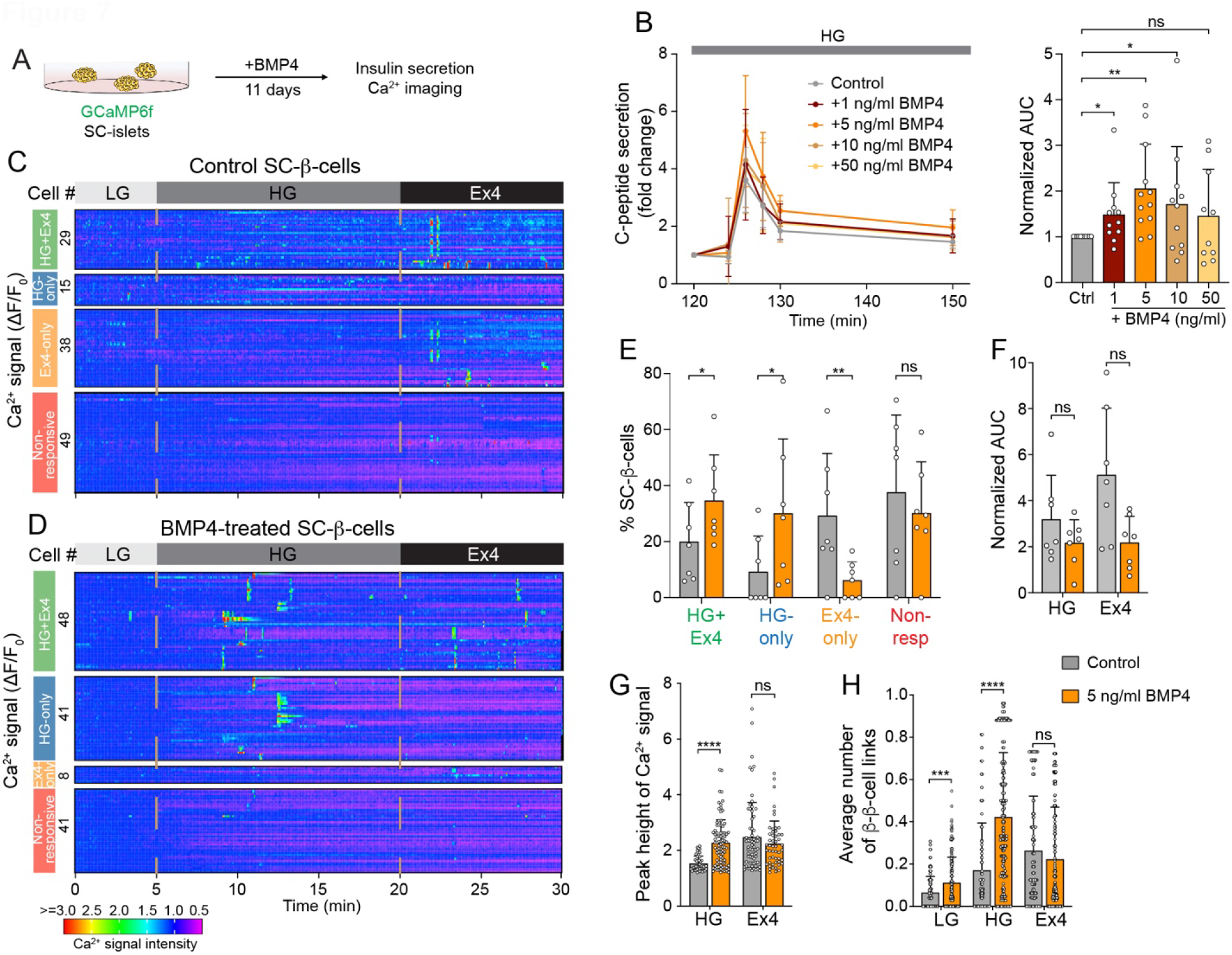
BMP4 treatment augments glucose-stimulated insulin secretion of SC-β-cells. (**A**) Schematic depicting workflow for BMP4 treatment and analysis of SC-islets. (**B**) C-peptide secretion (left) and quantification of area under the curve (AUC) (right) of control and BMP4-treated SC-islets during perifusion with 16.8 mM glucose (HG) (*n* = 10-12 pools of 50 SC-islets each; dots denote data points of individual pools). The AUC value in controls was set as 1. (**C** and **D**) Heatmaps showing normalized Ca^2+^ signals of 131 SC-β-cells of 7 control SC-islets and 138 SC-β-cells of 7 BMP4-treated SC-islets from 3 independent experiments exposed to LG, HG, and Ex4. High Ca^2+^ signal (red), low Ca^2+^ signal (violet). (**E**) Quantification of SC-β-cells exhibiting a Ca^2+^ response to HG and Ex4, HG but not Ex4 (HG-only), Ex4 but not HG (Ex4-only) or being non-responsive to either HG or Ex4 (Non-resp) in control and BMP4-treated SC-islets (*n* = 7 for both control and BMP4-treated SC-islets from 3 independent experiments). (**F**) Quantification of area under the curve (AUC) of SC-β-cells exhibiting a Ca^2+^ response in HG (from HG+Ex4 and HG-only groups) or Ex4 (from HG + Ex4 and Ex4-only groups). *n* = 7 for control and BMP4-treated SC-islets from 3 independent experiments; dots denote data points of individual SC-islets. The AUC value in LG was set as 1. (**G**) Quantification of Ca^2+^ signal peak height of responsive SC-β-cells in HG (from HG+Ex4 and HG-only groups; *n* = 44 SC-β-cells in control SC-islets and *n* = 89 SC-β-cells in BMP4-treated SC-islets) or Ex4 (from HG + Ex4 and Ex4-only groups; *n* = 67 SC-β-cells in control SC-islets and *n* = 56 SC-β-cells in BMP4-treated SC-islets; dots denote data points of individual SC-β-cells). (**H**) Quantification of connectivity of SC-β-cell Ca^2+^ response in SC-islets exposed to LG, HG, and Ex4 (*n* = 117 SC-β-cells in 7 control SC-islets and *n* = 193 SC-β-cells in 7 BMP4-treated SC-islets from 3 independent experiments; dots denote data points of individual SC-β-cells). All data are shown as mean ± S.D., Student’s *t* test, *p-value < 0.05, **p-value < 0.01, ***p-value < 0.001, ****p-value < 0.0001, ns, not significant.

## Discussion

In islets, β-cells are surrounded by a dense vascular network that enables β-cell glucose sensing and insulin secretion [14]. The proximity of vascular ECs to β-cells in the islet niche creates a local microenvironment that provides important cues for the regulation of β-cell function [15, 16]. Here, we describe the generation of a vascularized SC-islet organoid model comprised of SC-islets embedded into a 3D vascular network. Consistent with the importance of ECs for β-cell function, we show that the presence of vasculature improves the response of SC-β-cells to physiological stimuli of insulin secretion. Moreover, using single-cell analyses, we identify cell-cell communication networks between SC-β-cells and ECs that support formation of a functional organoid. Our demonstration of effects of ECM and EC-derived paracrine factors on SC-β-cells illustrates the value of the here-described vascularized SC-islet model as a tool for studying heterotypic cell-cell interactions in the islet microenvironment.

Studies in mice have shown that β-cells secrete VEGF-A which serves as an essential molecular cue for islet vascularization [14, 17, 30]. In the SC-islet organoid model, the same signaling mechanism from β-cells to ECs appears to be operating. The culture medium (GM1 medium) that supports both vascular network formation and SC-β-cell maintenance in the vascularized SC-islet organoid is devoid of VEGF-A in the last two days of the five-day culture period. After five days, we observed poor vascular network density when ECs and FBs were cultured in this medium without SC-islets, whereas intact vascular networks were present in co-culture with SC-islets. This suggests that SC-islets produce VEGF-A to support formation of a vascular network. Our cell-cell communication network analysis identified SC-β-cells as the likely source of VEGF-A signaling to ECs.

The cell-cell communication network also predicted reciprocal signaling between ECs and SC-β-cells. Consistent with ECs being the major source of islet basement membrane ECM proteins [17, 29, 41], vascularized SC-islet organoids contain basement membrane-like structures frequently contacted by SC-β-cells. We show that vascularization of SC-islets increases the percentage of SC-β-cells with active ECM-integrin-β1 signaling, which is known to stimulate β-cell insulin secretion [17, 19]. Accordingly, we found that culture of SC-islets in the presence of basement membrane-like ECM improves the SC-β-cell Ca^2+^ response to insulin secretory stimuli. These findings suggest that vasculature-derived ECM accounts for at least some of the observed functional improvement of SC-β-cells in the vascularized compared to the non-vascularized SC-islet model. Further demonstrating the importance of the islet microenvironment for proper β-cell function, ECM-induced integrin-β1 signaling at the side of the β-cell facing vasculature triggers a localized Ca^2+^ response and insulin exocytosis [18, 19, 42]. This local β-cell environment is not preserved in islets isolated from human pancreas, however, our vascularized SC-islet organoid now provides a model in which spatially localized signals from vasculature to β-cells can be studied. For example, there is predicted semaphorin signaling between ECs to SC-β-cells (Fig. 5C) which remains to be functionally explored.

The scRNA-seq analysis also revealed paracrine signals from ECs to SC-β-cells via soluble factors. We find that ECs are a source of BMP4 in vascularized SC-islet organoids and that BMP4 augments SC-β-cell function. Our findings are consistent with a recent study showing a role of pericyte-derived BMP4 in glucose-stimulated insulin secretion [43]. In addition, we show that BMP4 increases the magnitude of Ca^2+^ influx and synchronizes Ca^2+^ responses of neighboring SC-β-cells. Synchronicity of Ca^2+^ influx into β-cells is thought to underlie pulsatile insulin secretion which is a hallmark of mature islet function and perturbed during progression to type 2 diabetes [44, 45]. Thus, our findings suggest that BMP4 secreted by ECs could be an important maturation signal for β-cells. An additional candidate EC-derived signal acting on β-cells is progranulin which is endocytosed via the sortilin receptor. Progranulin regulates lysosomal function and stimulates β-cell proliferation in mouse islets [46]; however, its role in human β-cells remains to be studied. The compendium of candidate signals from ECs to SC-β-cells identified by our cell-cell communication network analysis provides a roadmap for further exploration of signaling mechanisms by which ECs influence β-cells.

One limitation of our vascularized SC-islet model is that it lacks the dense intra-islet vascular network characteristic of native islets. To enable formation of an intra-islet vascular network, we attempted to dissociate SC-islets into single cells or smaller clusters before aggregating with ECs and FBs. However, ECs self-aggregated or migrated out of the cell clusters, making it impossible to establish an intra-islet vascular network. Proper penetration of the organoid by vasculature is a known challenge in the organoid field, well-documented for liver and kidney organoids [47, 48]. Further analysis of our cell-cell communication data set is underway as we probe potential mechanisms underlying this phenomenon. Despite this limitation, our data show that the vasculature in our model is sufficient to significantly improve SC-β-cell function. Furthermore, in type 1 diabetes cytotoxic T cells accumulate in the peri-islet space, suggesting that T cell extravasation occurs from vessels outside and not inside the islet [49]. Therefore, the model described here could provide relevant insights into β-cell killing in type 1 diabetes.

Loss of islet function and disruption of vascular morphology are features of human type 1 [50] and type 2 diabetes [51]. In type 2 diabetes, these changes are thought to result from hyperglycemia [52], whereas cause and effect are less clear in type 1 diabetes [53]. Since vasculature is lost in primary islet cultures [22], the role of islet vasculature in disease cannot be studied in human islets. The microfluidic SC-islet model with perfused vasculature provides a new platform in which effects of diabetes-associated local and systemic environmental conditions can be studied. The flow through blood vessels in the microfluidic device permits mimicking humoral changes associated with diabetes *in vivo* to assess effects on vasculature and islet cells. Fluorescent probes, as here shown for the Ca^2+^ reporter GCaMP6f, can provide real time read-outs of cellular function and health in the device using live imaging. Given that stem cells are genetically tractable, the model further enables studies of how the cellular environment is interpreted in the context of specific genotypes.

Finally, the vascularized SC-islet organoid will serve as a platform to further engineer the cellular complexity of the pancreatic islet for disease modeling. Islet inflammation contributes to β-cell dysfunction and destruction and results from the interplay between local macrophages, adaptive immune cells, and vascular ECs [54, 55]. ECs are important for the recruitment of macrophages to the islet and homing of macrophages in the vascular niche [54, 56]. Signaling cascades between ECs, macrophages, and β-cells in the vascular niche initiate autoimmune insulitis in type 1 diabetes [57] and low-grade chronic inflammation in type 2 diabetes [58]. Like ECs, islet-resident macrophages and other immune cells are quickly depleted in cultured human islets, complicating *ex vivo* studies of immune-endocrine cell crosstalk. The SC-islet organoid model, with the presence of a functional vascular network and islet-like vascular niche, should facilitate the incorporation of immune cells to model mechanisms of β-cell dysfunction and destruction in diabetes. In the long-term, the goal is to develop a fully autologous SC-islet organoid model to study mechanisms of autoimmune β-cell destruction in type 1 diabetes.

## Acknowledgements

We acknowledge support of the Sanford Consortium for Regenerative Medicine Stem Cell Genomics and Microscopy Core, the UCSD Center for Epigenomics, the UCSD IGM Genomic Center, and the UCI Genomic High-Throughput Facility. This work was supported by grants from the Human Islet Research Network of the National Institutes of Health UC4DK104202 (M.Sa., C.C.W.H., S.C.G., K.L.K.), UG3/UH3DK122639 (M.Sa. and C.C.W.H.) and UC24 DK1041162 (V.K.), a grant from the Diabetes Research Connection (H.Z.), grant 25B1756 from the Burroughs Wellcome Fund (V.K.), and postdoctoral fellowships from the Juvenile Diabetes Research Foundation (K.V.N.N. and Y.J.) and the California Institute for Regenerative Medicine (H.Z.). Work at the Center for Epigenomics was supported in part by the UCSD School of Medicine. We thank Luc Teyton, Matthew Wortham, Bhupinder Shergill, Benjamen O’Donnell, Duc Phan, and members of the Sander lab for scientific discussions and input on the project.

## Author Contributions

M.Sa., C.C.W.H., K.L.C., S.C.G, and K.V.N.N. conceived the research. M.Sa., K.V.N.N., and Y.J. co-wrote the manuscript; K.V.N.N., and Y.J. designed and performed experiments and analyzed experimental data; K.V.N.N, S.S., C.J.H., H.Z., S.J.H., and M.M. generated and analyzed scRNA-seq data; V.K. analyzed cell-cell connectivity networks of Ca^2+^ signals; H.Z. generated the GCaMP6f-hESC line; R.G. contributed matrices while optimizing organoids; and M.Sch. contributed to Ca^2+^ imaging experiments and analyses. C.C.W.H. and R.H.F.B. advised on vascularization experiments, provided microfluidic devices, and contributed to experiments in devices.

## Materials and Methods

### Resource availability

#### Lead contact

Further information and requests for resources and reagents should be directed to and will be fulfilled by the Lead Contact, Maike Sander.

### Experimental model and subject details

#### Maintenance and SC-islet differentiation of H1 hESCs

All experiments with hESCs were approved by the University of California, San Diego (UCSD), Institutional Review Board and Embryonic Stem Cell Research Oversight Committee (protocol 090165ZX). H1 hESCs were maintained as described [59]. In brief, hESCs were seeded onto Matrigel (Corning, 356238) coated tissue culture surfaces in mTeSR1 media (Stem Cell Technologies, 85850) supplemented with 1% Penicillin-Streptomycin (Thermo Fisher Scientific, 15140122), and propagated every 3 to 4 days. Accutase (Thermo Fisher Scientific, 00-4555-56) based enzymatic dissociation method was employed for passaging and 10 µM Y-27632 (Stem Cell Technologies, 72307) was supplied on the first day of each passage.

H1 hESCs were differentiated into SC-islets with a protocol we modified from previous publications [12, 13]. After dissociation using Accutase, cells were reaggregated in 5.5 mL mTeSR1 supplemented with Y-27632 (Stem Cell Technologies) by suspending the cells at a concentration of 5.5 × 10^6^ cells/well in a low attachment 6-well plate on an orbital shaker (100 rpm) in a 37°C incubator. The following day (day 0), undifferentiated cells were washed in Stage 1/2 base medium (see below) and then differentiated using a seven-step protocol with stage-specific medium. Medium was refreshed daily until day 32 and every two days thereafter. On day 21, cells were dissociated with Accutase, suspended in Stage 7 medium (see below) supplemented with 10 µM Y-27632 and re-aggregated at a concentration of 3 × 10^6^ cells/well in a low attachment 6-well plate on an orbital shaker (100 rpm) in a 37 °C incubator. The speed of the shaker was increased to 118 rpm on the following day. Co-culture studies with ECs and FBs were initiated from SC-islets > 39 days. The differentiation protocol and media composition are detailed in **Table S4**.

#### Generation of H1 GCaMP6f-hESCs

The GCaMP6f-hESC line was generated by knock-in of GCaMP6f into the human *AAVS1* locus [60]. In brief, we cloned the sgRNA targeting *AAVS1* locus, sgAAVS1, (GGGGCCACTAGGGACAGGAT) [61] into a Cas9-expressing vector, pX458 (Addgene #48138). H1 hESCs were then co-transfected with sgAAVS1-pX458 and a GCaMP6f donor vector (Addgene #73503) containing *AAVS1*-targeting recombination arms [62] using XtremeGene 9 (Roche). Transfected cells were allowed to recover for 48 hrs before treatment with 1 μg/ml puromycin (Sigma) for an additional 48 hrs. 72 hrs after puromycin removal, 8000 cells were plated into a well of 6-well plate. Individual colonies that emerged within 5–7 days were subsequently transferred manually into 48-well plates for expansion, genomic DNA extraction, PCR genotyping, and Sanger sequencing. The following primers were used for genotyping: AAVS1 upstream forward: CTGCCGTCTCTCTCCTGAGT AAVS1 upstream reverse: GTGGGCTTGTACTCGGTCAT AAVS1 downstream forward: TCCCAGTCATAGCTGTCCCTCT AAVS1 downstream reverse: TGGTAGACAGGGCTGGGGTG.

#### Primary islet culture

Human cadaveric islets were acquired through the Integrated Islet Distribution Program (IIDP). Upon arrival, islets were stained with dithizone and stained islets hand-picked under a dissection microscope. Islets were allowed to recover overnight in islet medium (CMRL 1066 medium with 13 mM D-Glucose, 10% FBS, 2 mM L-glutamine, 100U/ml Penicillin-Streptomycin, 1 mM sodium pyruvate, 10 mM HEPES, 0.25 µg/ml Amphotericin B).

#### Culture of primary endothelial cells (ECs) and fibroblasts (FBs)

For primary ECs, human umbilical vein endothelial cells (HUVEC) were acquired from Lonza and cultured in EGM-2 medium (Lonza) on a 0.1% gelatin-coated tissue culture dish. EGM-2 medium (referred as vascularization medium, VM) contains endothelial basal medium (EBM) and the following growth factor supplements: human Epidermal Growth Factor (EGF), Vascular Endothelial Growth Factor (VEGF), Insulin-like Growth Factor-1 (IGF), Ascorbic acid, Hydrocortisone, human basic Fibroblast Growth Factor (bFGF), Heparin, Fetal Bovine Serum (FBS), Gentamicin/Amphotericin-B (GA). At passage 2, ECs were transduced with lentivirus expressing fluorescent proteins (Azurite Blue) for network visualization. For primary FBs, normal human lung fibroblasts (NHLF) were acquired from Lonza and cultured in an FGM-2 BulletKit (Lonza) on a tissue culture dish. Both ECs and FBs were passaged by splitting at 80% confluency using 0.25% trypsin/EDTA (Thermo Fisher Scientific) and used for co-culture study between passages 4-8.

#### Generation of vascularized SC-islet organoids in 3D fibrin gels

To create uniformly sized SC-islet clusters (< 200 µm in diameter), SC-islets were dissociated into single cells using Accutase for 10 min and seeded into a BSA-coated concave-shaped microwell platform (StemFIT 3D, microFIT) at a density of ∼ 1000 cells per microwell (diameter, 400 µm) using the manufacturer’s protocol. SC-islets were allowed to reaggregate for at least 2 days before setting up co-cultures with ECs and FBs. To prepare 3D fibrin gels, fibrinogen powder (Sigma-Aldrich) was dissolved in EBM medium to a final concentration of 2 mg/ml before filtering. 100 resized SC-islets collected from the microwells and 2 × 10^5^ dissociated ECs and FBs (1:1) were resuspended in 100 μl fibrinogen before mixing cells with 16 μl of 50 U/ml thrombin (Sigma-Aldrich). The mixture was plated into one well of a 24-well plate to create a round-shaped gel droplet and allowed to polymerize at 37 °C for 30 min. Upon gelation, each fibrin gel was filled with different co-culture media described in **Table S1**. The co-cultures were maintained for 4-7 days. Vascular networks and SC-islet morphology were examined daily by bright-field and epi-fluorescent microscopy. Non-vascularized aggregates that include only resized SC-islets were also embedded and cultured in a fibrin gel for the same period.

#### Incorporation of vascularized SC-islet organoids into microfluidic devices

Microfluidic device used in the study was fabricated using standard soft lithography techniques as previously described [26]. The PDMS (Sylgard 184, Dow Corning) device consists of two straight microfluidic channels (upper channel as arteriole and lower channel as venule) separated by a tissue chamber (channel height, 200 µm). The channels are connected to two medium inlets and outlets on each side. Medium reservoirs are attached to the device using PDMS. All devices were subsequently sterilized by autoclaving at 120 °C before use. We followed the method for loading cells into microfluidic channels and forming vessels as previously described [26]. Briefly, to prepare fibrin gels for device loading, the filtered fibrinogen solution in EBM at 5 mg/ml was supplemented with 0.15 U/ml Aprotinin (Sigma-Aldrich) at the ratio of 1:100 (v/v), then mixed with 0.2 mg/ml fibronectin at the ratio of 1:4 (v/v). Next, 100-130 resized SC-islets from the microwells 35 × 10^3^ dissociated ECs and FBs (1:1) were thoroughly resuspended in 5 μl of fibrinogen-fibronectin solution, quickly mixed with 1.2 μl of 50 U/ml thrombin (Sigma-Aldrich) before loaded into a central tissue chamber. After incubation at 37 °C for 10 min, the device was taken out to have the upper and lower channels coated with 3.5 μl of fibronectin/each followed by an additional 15-minute incubation. Finally, medium was added to the reservoirs to prevent bubble formation. The volume of medium in four reservoirs was alternated every two days to induce vessel formation and anastomosis with both channels. At day 5, 1 ml of 70 kDa fluorescein-dextran (green)-containing medium was perfused through the tissue chamber to test vessel leakiness. The perfused co-culture was maintained in the device for up to 7 days using the culture conditions described in **Table S2**.

#### Immunofluorescence staining and image analysis

SC-islets, cadaveric islets, and tissues in fibrin gels were washed twice with PBS and fixed with 4% paraformaldehyde (PFA) for 30 min at room temperature (RT). Fixed samples were washed twice with PBS and incubated overnight at 4°C in 30% (w/v) sucrose solution (Thermo Fisher Scientific) in PBS. Samples were then embedded into disposable embedding molds (VWR), covered in Tissue-Tek® O.C.T. Sakura® Finetek compound (VWR), flash frozen on dry ice, and stored at -80°C. Tissue blocks were sectioned at 10 μm and sections were placed on Superfrost Plus® (Thermo Fisher) microscope slides. For immunofluorescence staining, slides were thawed at RT, washed twice with PBS for 10 min, permeabilized with 0.15% (v/v) Triton X-100 (Sigma-Aldrich) for 30 min, and blocked with blocking buffer, consisting of 0.15% (v/v) Triton X-100 (Sigma-Aldrich) and 1% (v/v) normal donkey serum (Jackson Immuno Research Laboratories) in PBS for 1h at RT. Slides were then incubated overnight at 4°C with primary antibodies diluted in blocking buffer. The following day slides were washed thrice with blocking buffer for 15 min and incubated for 1 h at RT in secondary antibodies and Hoechst 33342 (Invitrogen) diluted in blocking buffer. Stained sections were washed three times with PBS before mounting with VECTASHIELD^®^ (Vector Laboratories). Images were obtained with a Zeiss Axio-Observer-Z1 microscope equipped with a Zeiss ApoTome and AxioCam digital camera. For samples from the device, tissue channels were perfused with 4% PFA for 30 min at RT, then flushed with PBS before the bottom silicon membrane was removed. The tissue was either stained as a whole or cryosectioned. Cell populations were quantified and analyzed using HALO image analysis platform (Indica Lab). One section was analyzed for each SC-islet.

#### Dynamic insulin secretion assay and C-peptide measurements

Dynamic insulin secretion assays were carried out using the Biorep perifusion system (Biorep v5) with a flow rate of 100 µl/min, in the Krebs-Ringers-Bicarbonate-HEPES (KRBH) buffer that contains 130 mM NaCl, 5 mM KCl, 1.2 mM CaCl_2_, 1.2 mM MgCl_2_, 1.2 mM KH_2_PO_4_, 20 mM HEPES pH 7.4, 25 mM NaHCO_3_, and 0.1% BSA. 30-50 hand-picked SC-islets were loaded into the perifusion chamber and equilibrated in KRBH buffer with 2.8 mM glucose for 2h, followed by 16.8 mM glucose for 45 min, 16.8 mM glucose with 10 nM Exendin-4 for 15 min, 2.8 mM glucose for 1h, and 30 mM KCl for 20 min. Samples were collected and analyzed using STELLUX® Chemi Human C-peptide ELISA (ALPCO). SC-islets from perifusion chambers were transferred to microcentrifuge tubes and lysed by sonication for total C-peptide content measurement. Total DNA was measured by Nanodrop spectrophotometer (Thermo Fisher Scientific).

For analysis of vascularized SC-islets, SC-islet-containing gels were dislodged, collected into microcentrifuge tubes, and lysed by sonication for 10 cycles before proceeding to total C-peptide and DNA content measurements. Total C-peptide content was normalized to total DNA.

#### Ca^2+^ imaging and analysis

Fibrin-embedded GCaMP6f-SC-islets were placed onto the glass-bottom 4-chamber slide (Thermo Fisher). After washing SC-islets with KRBH, they were live stained with anti-CD49a-PE antibody in KRBH contained 2.8 mM glucose for 1h at 37°C. During the last 5 min of the 1h incubation, Hoechst 33342 was added for 5 min before washing three times with KRBH containing 2.8 mM glucose and adding 500 μl of fresh KRBH with 2.8 mM glucose. A still image of CD49a, DAPI, and GFP was taken right before recording the Ca^2+^ signal every 5s for 5 min at 2.8 mM glucose, 10-15 min at 16.8 mM glucose, and 10 min at 16.8 mM glucose + 10 nM Exendin-4. Afterward, the fibrin-embedded SC-islets were fixed and stained with anti-insulin, anti-NKX6.1, and anti-CD49a-PE antibodies. Confocal images were taken in the same focal plane as the previously recorded Ca^2+^ signals in the SC-islet. All images were taken using a 40x water objective of a Zeiss 780 microscope.

A similar workflow was applied for Ca^2+^ imaging of SC-islets in the microfluidic device. In the device, SC-islets were perfused with KRBH buffer solutions and imaged using a 20x objective of a Zeiss 780 microscope.

Intracellular Ca^2+^ signals of SC-β-cells were analyzed by superimposing upon each other (i) live images of CD49a-PE, GFP, and DAPI, (ii) images of immunofluorescent staining, and (iii) time-lapse images of Ca^2+^ signals. SC-β-cells were selected based on expression of insulin, NKX6.1, and CD49a. Background-subtracted intensity of GCaMP6f fluorescence (ΔF) was used to calculate ΔF/F_0_ by normalizing ΔF values to the mean values of ΔF at 2.8 mM glucose (5 min, 60 cycles) (F_0_). A cell was defined as ‘responsive’ if it was elicited at least twice under the same stimulus with the amplitude of the peak being greater than 1.2-fold of the baseline fluctuation (ΔF/F_0_ ≥ 1.2). Based on this criterion, SC-β-cells were categorized into four groups depending on their responsiveness: (i) responsive to both high glucose (16.8 mM) and Exendin-4, (ii) responsive to only high glucose but not Exendin-4, (iii) responsive to Exendin-4 but not high glucose, and (iv) not response to either high glucose or Exendin-4. The area under the curve (AUC) of individual Ca^2+^ traces of responsive SC-β-cells was quantified for each stimulus. All calculations were made using ImageJ (National Institute of Health) and Prism (GraphPad) software.

#### Network analysis of Ca^2+^ dynamics

Pearson-product-based network analysis was performed as previously reported [40]. Ca^2+^ signals were analyzed during exposure to low glucose, high glucose, and Exendin-4, where 5 min intervals for each of the conditions was chosen such that the cluster exhibited highest activity. The Pearson product for each cell pair in the SC-islet was calculated over each time point, and the time-average values were computed to construct a correlation matrix. An adjacency matrix was calculated by applying a threshold of 0.30 to the correlation matrix. The same threshold was applied to all clusters and conditions. All cell pairs with non-zero values in the adjacency matrix were considered to have a functional link. The average number of links per cluster was calculated as mathematical average of all non-zero links in the cluster. It was then normalized by the number of cells in the given cluster to take into account differences in cluster sizes.

#### Single-cell RNA sequencing library preparation

SC-islet cells at day 32 and 39 of differentiation were collected and washed with PBS. Cell aggregates were dissociated into a single-cell suspension with Accutase at 37°C. Cells were then resuspended into 1 ml of ice-cold buffer consisting of 0.2% BSA in PBS and stained with propidium iodide (Sigma, Cat# P4170) to eliminate dead cells. 500,000 live cells were collected using a FACSAria™ Fusion Flow Sorter, and 10,000 cells per sample were then loaded onto a 10X Chromium Controller and run using Next GEM Single-Cell 3’ v3.1 reagents. Library was prepared according to manufacturer’s instructions and sequenced using a NovaSeq S4 (Paired-end 100 bp reads, Illumina) at UCSD Genomics Facility.

ECs and FBs were retrieved from the tissue chamber of the microfluidic device after the thin transparent membrane was carefully removed. The chambers were washed with HBSS before adding TrypLE express enzyme in a drop-wise manner. After incubation at 37°C for 5 min, the chambers were flushed with a micropipette to collect 10,000 cells into a centrifuge tube. After quenching with EGM-2, cells were centrifuged (340xg for 3 min) and resuspended in an HBSS/collagenase type III solution (1 mg/mL) to digest the matrix completely. Dissociated cells were washed with EGM-2 and pushed through a 70 μm filter to create a single-cell suspension before being stained with propidium iodide to eliminate dead cells and sorted with a FACSAriaII. Finally, cells were loaded onto 10X Genomics Chromium and run using Chromium Single Cell 3’ v2 reagents. Library was prepared according to manufacturer’s instructions and sequenced using a HiSeq4000 at UCI Genomics High Throughput Facility (Paired-end 100 bp reads, Illumina).

#### Single-cell RNA sequencing analysis

Sequencing reads alignment and quantification were performed using 10X Cellranger (version 3.1.0) [63] with recent human genome-based reference from refdata-cellranger-GRCh38-3.0.0. The matrices generated by CellRanger were imported into Seurat (version 3.1.2 for SC-islets [64] and 4.0.0 for ECs and FBs [65]) for downstream processing. Cells were optimized based on doublet score < 0.3 and cells (>7500 total features for SC-islet day 32 and >6500 total features for SC-islet day 39 cells), low-coverage cells (<1500 total features for day 32 cells and < 1000 total features for day 39 cells), ribosomal genes and poor-quality cells (>20% mitochondrial reads for SC-islets, >10% for ECs and FBs) were removed before further analysis. Each dataset was Log Normalized with a scale factor of 10,000 using the command “NormalizeData”. Percentage of mitochondrial genes were regressed out of each dataset using the command “ScaleData”. Principal component analysis was performed for the using the command “RunPCA”, clusters were defined by running the commands “FindNeighbors” using 1:15 dimensions of reduction and “FindClusters” at a resolution of 0.02. The UMAP plots were generated through “RunUMAP” and marker genes were identified using “FindMarkers”.

SC-islet datasets and datasets from the microfluidic devices were integrated using Seurat’s SCT integration [64]. Briefly, each dataset was processed using Seurat’s SCTransform default settings. The top 3000 key features for integration were identified by running SelectIntegrationFeatures. To prepare for integration, the PrepSCTIntegration function was run. Then, the FindIntegrationAnchors and IntegrateData functions were used with their default settings to integrate the datasets. After integration, data were scaled before running a principal component analysis and UMAP before identifying clusters.

#### Analysis of cell-cell communication networks using CellChat

To infer cell-cell communication networks between SC-β-cells, FBs and ECs, CellChat [28] R package was used. The Seurat object, containing only the cell types of interest, was imported into CellChat via *createCellChat()* function and then processed using the standard pipeline. *CellChatDB*.*human* was used as the ligand-receptor database. It contains 1939 validated molecular interactions. Overexpressed ligands or receptors were identified in each cell-type with *identifyOverExpressedGenes()* and further interactions were identified with *identifyOverExpressedInteractions()*. Biologically significant cell-cell communication networks were inferred using *computeCommunProb()* function with parameters *type = “truncatedMean”* and *trim* set to 0.1. For further downstream analysis, self-pairs such as SC-β:SC-β, VMO-FBs:VMO-FBs and VMO-ECs:VMO-ECs were disregarded.

#### Quantification and statistical analysis

All experiments were repeated at least three times, and the data are presented as the mean ± standard deviation (SD). Statistical analyses were performed using GraphPad Prism (v8.1.2). The significance of group differences was evaluated with a two-tailed Student’s *t* test or analysis of variance (ANOVA).

**Figure S1.**
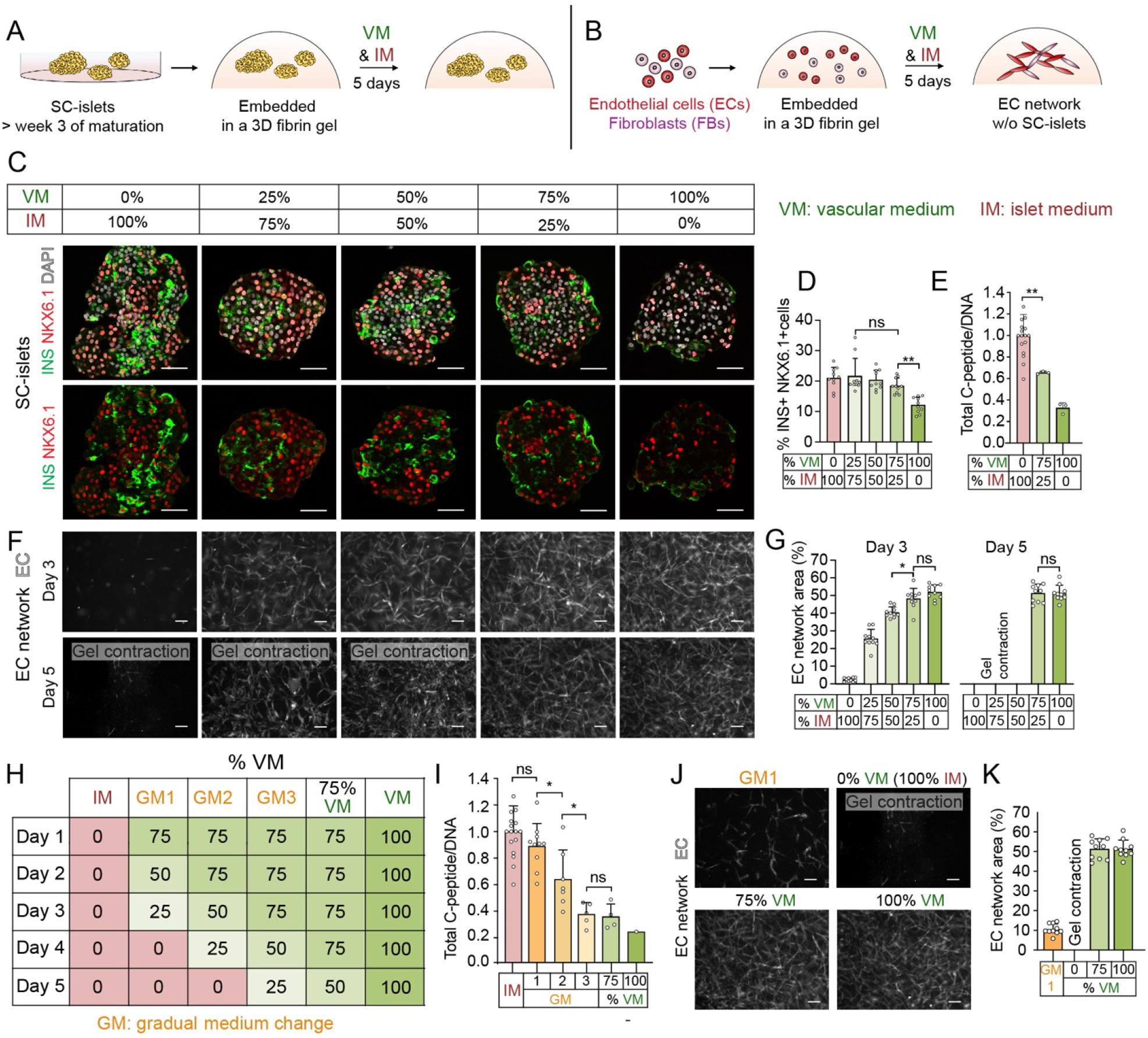
Optimization of culture conditions for vascularizing SC-islets. (**A** and **B**) Schematic depicting the method to test the effect of mixed VM and IM on SC-islets (A) and vascular network formation from ECs and FBs (B). (**C**) Representative immunofluorescence images of SC-islets stained with insulin (INS), NKX6.1, and DAPI after culture in different VM and IM mixtures for 5 days. Scale bars, 50 μm. (**D**) Quantification of SC-β-cells (NKX6.1+, INS+) in SC-islets after culture in different VM and IM mixtures for 5 days (*n* = 10 from 3 independent experiments). Dots denote data points of individual SC-islets. (**E**) Quantification of C-peptide content in SC-islets after culture in different VM and IM mixtures for 5 days (*n* = 17 for 100% IM from 10 independent experiments and *n* = 3 for two other conditions from 3 independent experiments). Dots denote data points of individual 3D fibrin gels containing 50 SC-islets each. (**F**) Representative epifluorescence images of vascular networks after culture in different VM and IM mixtures for 3 or 5 days. ECs were transduced with an azurite blue-expressing lentivirus. Scale bars, 50 μm. (**G**) Quantification of vascular network density at day 3 and 5 of culture in different VM and IM mixtures (*n* = 10 from 3 independent experiments). Dots denote data points of individual 3D fibrin gels. (**H**) Composition of different media tested. The percentage of VM relative to IM is indicated. GM, gradual medium changes. (**I**) Quantification of C-peptide content in SC-islets cultured under conditions shown in (H) for 5 days (*n* = 17 for IM, *n* = 10 for GM1, *n* = 7 for GM2, *n* = 5 for GM3, *n* = 4 for 75% VM, and *n* = 1 for 100% VM from 10, 6, 5, 4, 3 and 1 independent experiments, respectively). Dots denote data points of individual 3D fibrin gels containing 50 SC-islets each. (**J**) Representative epifluorescence images of EC networks cultured under conditions shown in (H) for 5 days. ECs were transduced with an azurite blue-expressing lentivirus. Scale bars, 50 μm. (**K**) Quantification of vascular network density after culturing ECs and FBs for 5 days under conditions shown in (H) (*n* = 10 from 3 independent experiments; dots denote data points of individual 3D fibrin gels). All data are shown as mean ± S.D., Student’s *t* test, *p-value < 0.05, **p-value < 0.01, ns, not significant.

**Figure S2.**
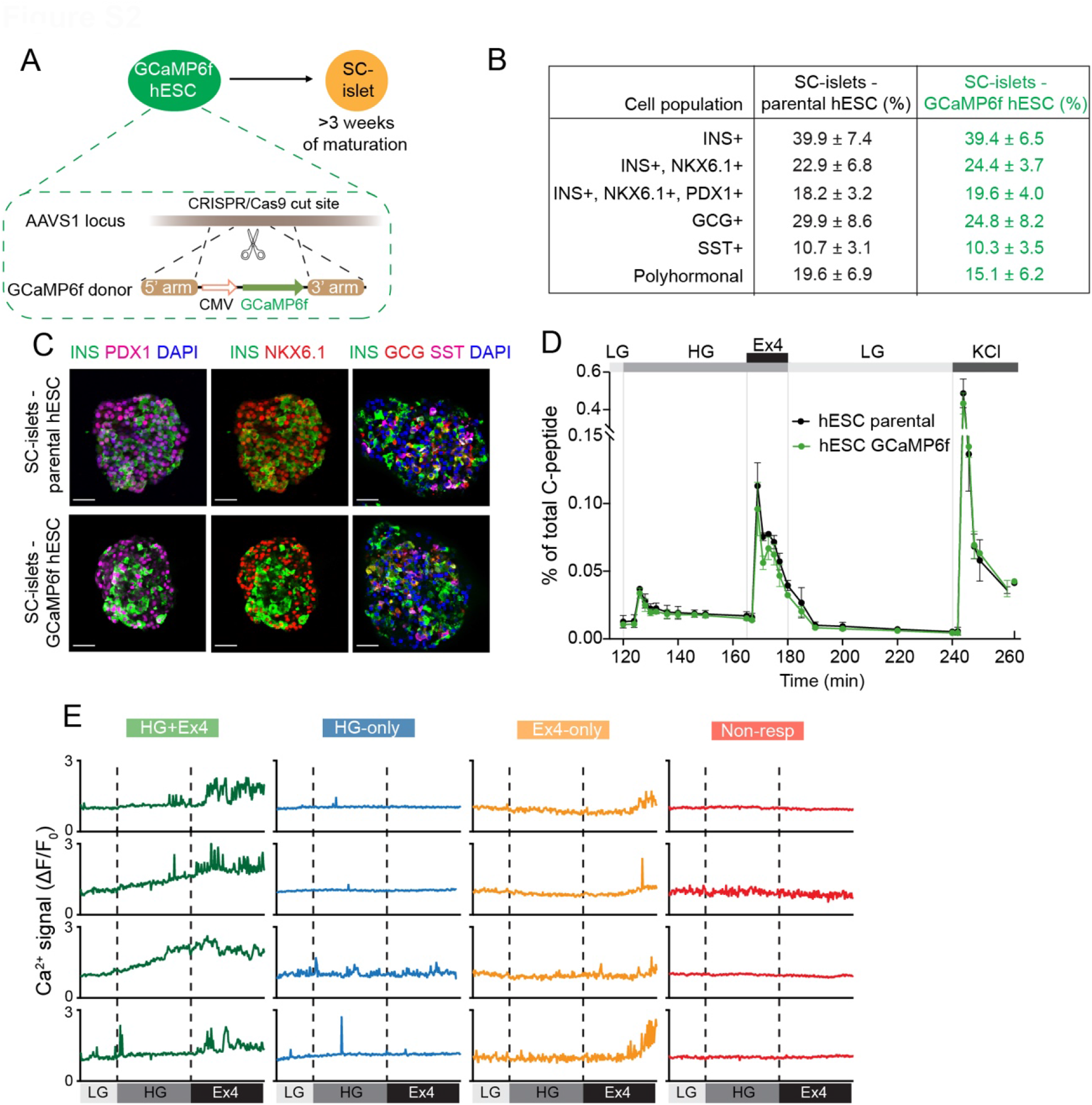
Characterization of GCaMP6f-SC-islets. (**A**) Schematic depicting generation and differentiation of GCaMP6f-hESCs. (**B**) Percentages of hormone-producing cells in SC-islets differentiated from parental hESCs and GCaMP6f-hESCs. Cell numbers relative to the total cell number in SC-islets quantified based on immunostaining for indicated markers (*n* = 10 SC-islets from 3 independent experiments; data are shown as mean ± S.D.). (**C**) Representative immunofluorescence images of SC-islets stained with the indicated markers. INS, insulin; GGC, glucagon; SST, somatostatin. Scale bars, 50 μm. (**D**) C-peptide secretion by SC-islets differentiated from parental hESCs and GCaMP6f-hESCs during perifusion with 2.8 mM glucose (LG), 16.8 mM glucose (HG), 10 nM Exendin-4 (Ex4), and 30 mM KCl (*n* = 3 pools of 50 SC-islets each). (**E**) Representative Ca^2+^ traces of SC-β-cells responsive to HG and Ex4, HG but not Ex4 (HG-only), Ex4 but not HG (Ex4-only) or not responsive to either HG or Ex4 (Non-resp).

**Figure S3.**
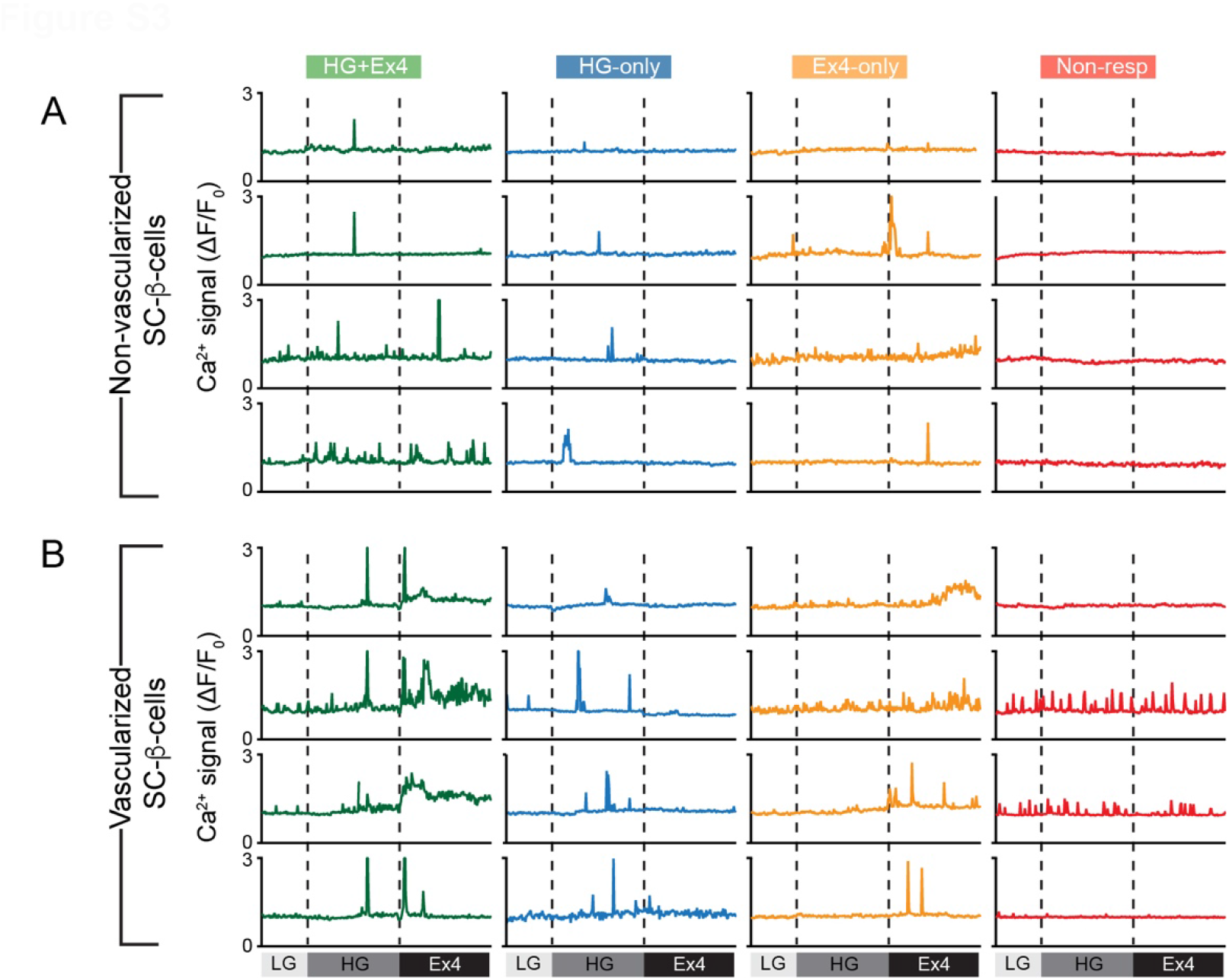
Ca^2+^ response of SC-β-cells in non-vascularized and vascularized SC-islet organoids. (**A** and **B**) Representative Ca^2+^ traces of SC-β-cells responsive to HG and Ex4, HG but not Ex4 (HG-only), Ex4 but not HG (Ex4-only) or not responsive to either HG or Ex4 (Non-resp) in non-vascularized (A) and vascularized (B) SC-islet organoids.

**Figure S4.**
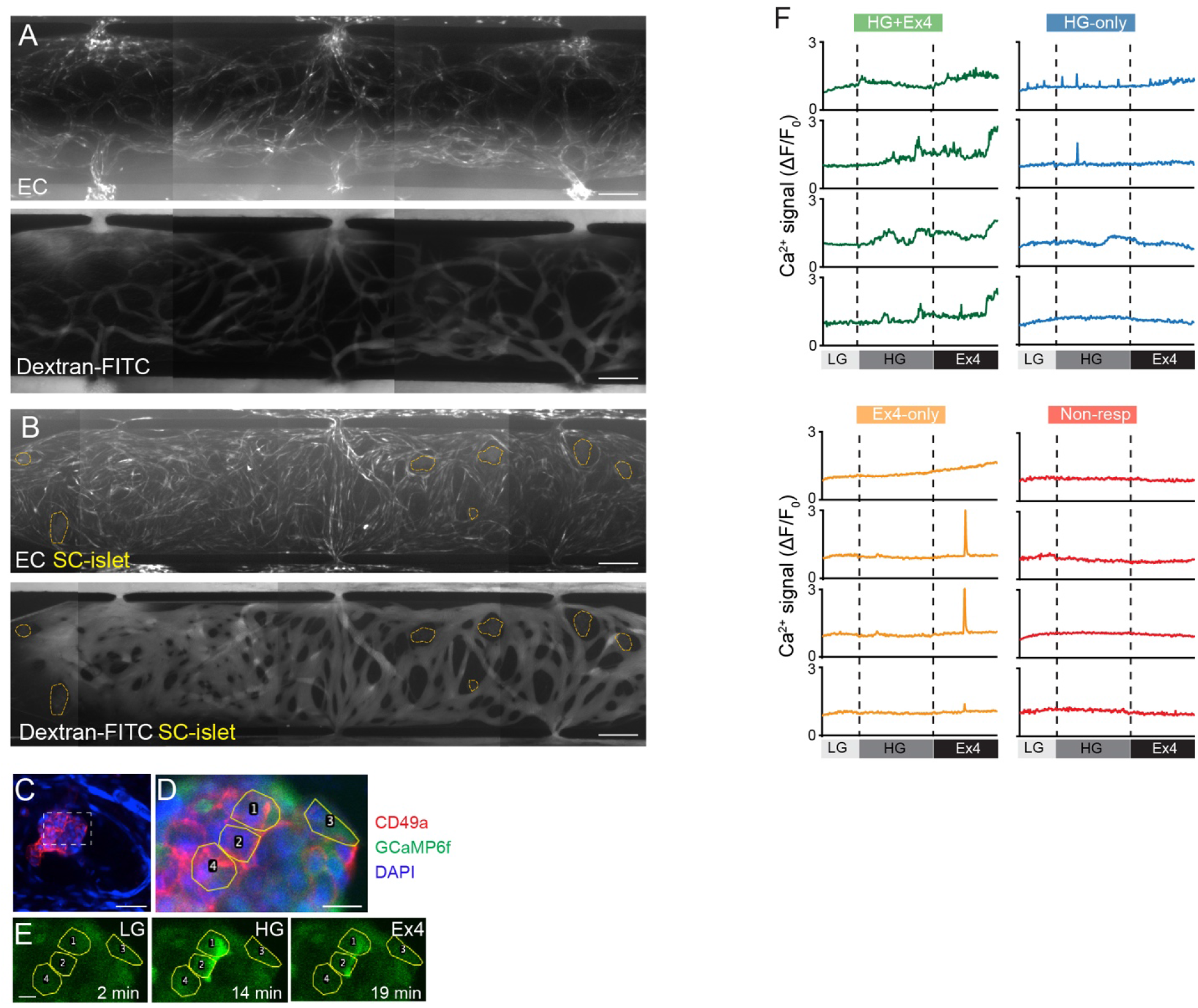
SC-islets are supported by a perfusable vascular network in the microfluidic device. (**A**) Representative epifluorescence images of live vascular network (top) and vascular network perfused with Dextran-FITC on day 6 in the device cultured in 100% VM without SC-islets. ECs were transduced with an azurite blue-expressing lentivirus. Scale bars, 200 μm. (**B**) Representative epifluorescence images of live vascular network (top) and vascular network perfused with Dextran-FITC in vascularized SC-islet organoids on day 6 in the device cultured in GM1 condition. SC-islets are circled. Scale bars, 200 μm. (**C** and **D**) Live image of a vascularized SC-islet organoid in the device stained with CD49a and DAPI. GCaMP6f signal is shown in green. Inset in C is shown in D with SC-β-cells selected for Ca^2+^ signal analysis. Scale bars, 50 μm (C) and 10 μm (D). (**E**) Still images of Ca^2+^ signals of select SC-β-cells sequentially exposed to 2.8 mM glucose (LG), 16.8 mM glucose (HG), and 10 nM Exendin-4 (Ex4). (**F**) Representative Ca^2+^ traces of SC-β-cells responsive to HG and Ex4, HG but not Ex4 (HG-only), Ex4 but not HG (Ex4-only) or not responsive to either HG or Ex4 (Non-resp) in the device.

**Figure S5.**
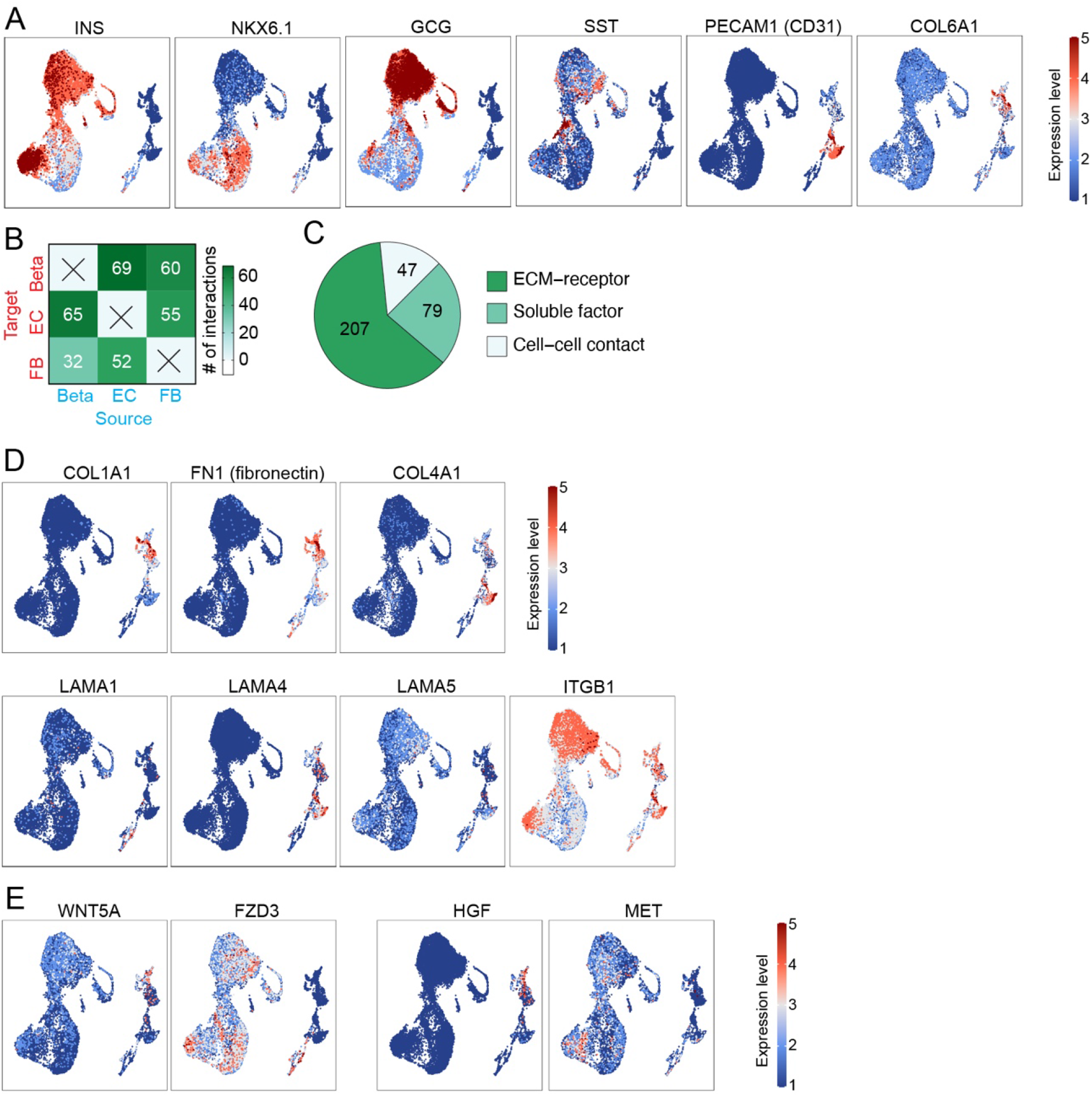
Predicting cell-cell communication networks between SC-β-cells, ECs, and FBs using scRNA-seq data. (**A**) Feature plots showing expression levels of marker genes for SC-islet cell types, ECs, and FBs. INS, insulin; GCG, glucagon; SST, somatostatin. (**B**) Number of predicted interactions between SC-β-cells, ECs, and FBs. (**C**) Distribution of predicted cell-cell interactions from B. (**D** and **E**) Feature plots showing expression levels of extracellular matrix proteins and integrin-β1 (ITGB1) (D), WNT5A and HGF ligands and their receptors in SC-islet cells, ECs, and FBs (E). High expression (red) and low expression (blue).

**Figure S6.**
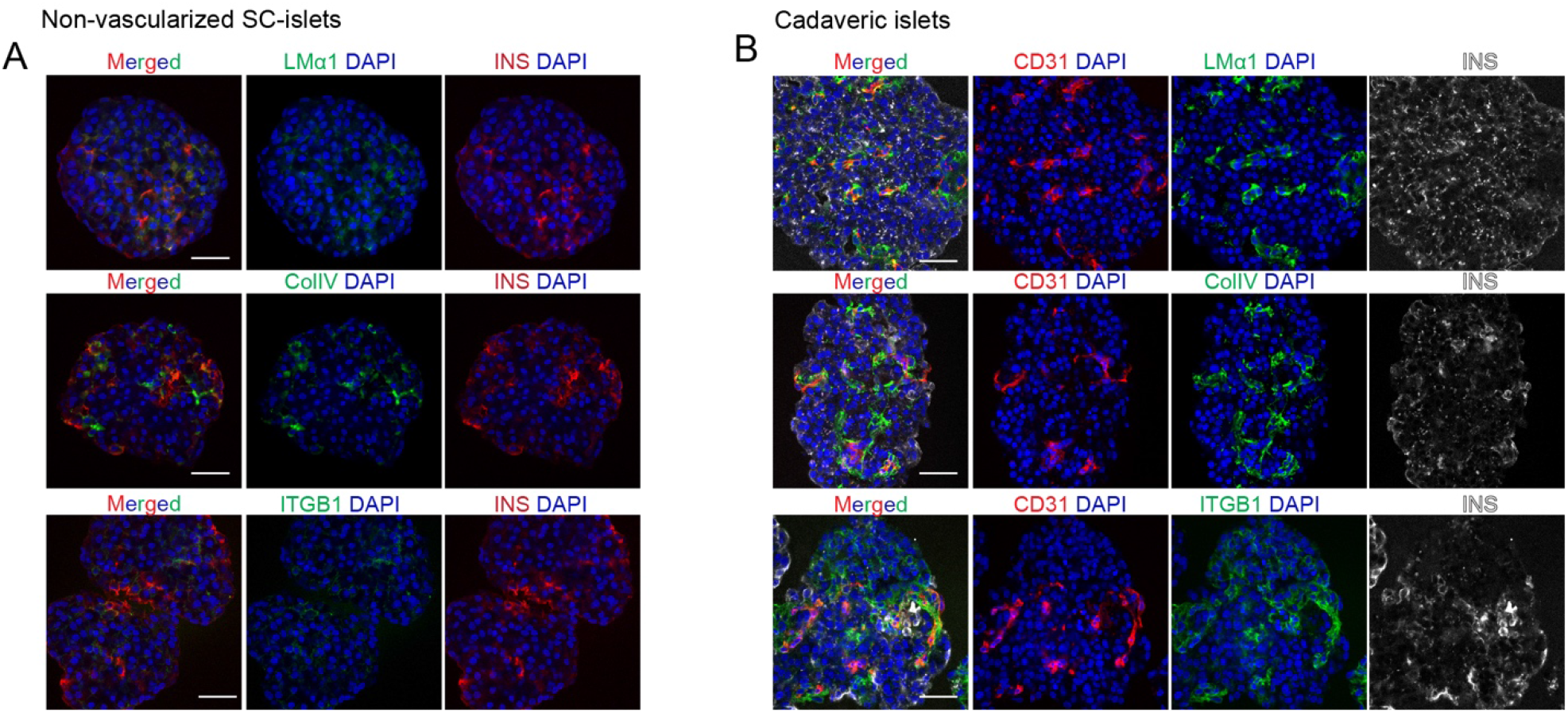
Non-vascularized SC-islets and cadaveric islets are devoid of a peri-basement membrane. (**A** and **B**) Representative immunofluorescence images of non-vascularized SC-islets (A) and cadaveric human islets (B) stained with laminin-α1 (LMα1), collagen IV (ColIV), integrin-β1 (ITGB1), CD31, and insulin (INS). Scale bars, 50 μm.

**Figure S7.**
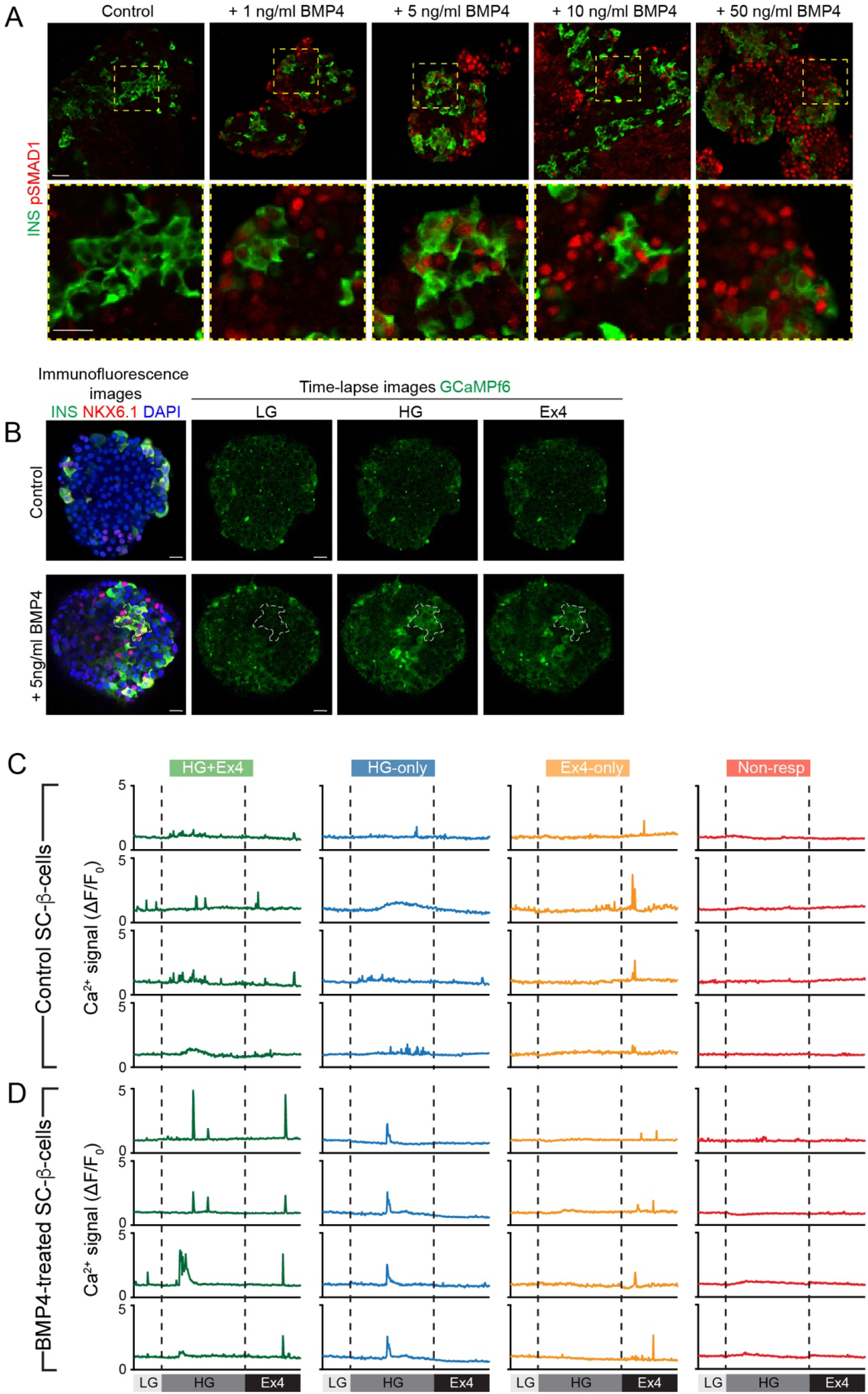
Effects of BMP4 treatment on SC-β-cells. (**A**) Representative immunofluorescence images of control and BMP4-treated SC-islets stained with insulin (INS) and pSMAD1. Bottom row shows magnifications of boxed areas in top row. Scale bars, 50 μm. (**B**) Representative immunofluorescence images stained for insulin (INS), NKX6.1, and DAPI (left) and still images of Ca^2+^ signals (right) in control and BMP4-treated SC-islets exposed to LG, HG, and Ex4. Scale bars, 20 μm. White dash-circled areas highlight a group of SC-β-cells showing Ca^2+^ response to HG and Ex4. (**C** and **D**) Representative Ca^2+^ traces of SC-β-cells responsive to HG and Ex4, HG but not Ex4 (HG-only), Ex4 but not HG (Ex4-only) or not responsive to either HG or Ex4 (Non-resp) in control (C) and BMP4-treated (D) SC-islets.

## Supplemental Information

**Video S1. Time-lapse video of live-imaged GCaMP6f-SC-islet exposed to low glucose (5 minutes), high glucose (15 minutes) and Exendin-4 (10 minutes)**. GFP fluorescent signals show Ca^2+^ influx into individual cells in GCaMP6f-SC-islets. Images were taken every 5 seconds for 30 minutes using 40x water objective. Video corresponds to Fig. 2D, relating to Fig. S2E.

**Video S2. Time-lapse video of live-imaged non-vascularized GCaMP6f-SC-islet cultured in GM1 for 5 days and exposed to low glucose (5 minutes), high glucose (10 minutes) and Exendin-4 (10 minutes)**. GFP fluorescent signals show Ca^2+^ influx into individual cells in GCaMP6f-SC-islets. Images were taken every 5 seconds for 25 minutes using 40x water objective. Video corresponds to Fig. 3B, relating to Fig. S3A.

**Video S3. Time-lapse video of live-imaged vascularized GCaMP6f-SC-islet cultured in GM1 for 5 days and exposed to low glucose (5 minutes), high glucose (10 minutes) and Exendin-4 (10 minutes)**. GFP fluorescent signals show Ca^2+^ influx into individual cells in GCaMP6f-SC-islets. Images were taken every 5 seconds for 25 minutes using 40x water objective. Video corresponds to Fig. 3C, relating to Fig. S3B.

**Video S4. Time-lapse video of live-imaged vascularized GCaMP6f-SC-islet cultured in microfluidic device for 6 days and exposed to low glucose (5 minutes), high glucose (13 minutes) and Exendin-4 (10 minutes)**. GFP fluorescent signals show Ca^2+^ influx into individual cells in GCaMP6f-SC-islets. Images were taken every 5 seconds for 25 minutes using 20x water objective. Video corresponds to Fig. 4D, relating to Fig. S4F.

**Video S5. Time-lapse video of live-imaged GCaMP6f-SC-islet encapsulated into a 3D fibrin gel and exposed to low glucose (5 minutes), high glucose (15 minutes) and Exendin-4 (10 minutes)**. GFP fluorescent signals show Ca^2+^ influx into individual cells in GCaMP6f-SC-islets. Images were taken every 5 seconds for 30 minutes using 40x water objective. Video corresponds to Fig. 6H.

**Video S6. Time-lapse video of live-imaged GCaMP6f-SC-islet encapsulated into a 3D fibrin+Matrigel (1:1) gel and exposed to low glucose (5 minutes), high glucose (15 minutes) and Exendin-4 (10 minutes)**. GFP fluorescent signals show Ca^2+^ influx into individual cells in GCaMP6f-SC-islets. Images were taken every 5 seconds for 30 minutes using 40x water objective. Video corresponds to Fig. 6I.

**Video S7. Time-lapse video of live-imaged GCaMP6f-SC-islet exposed to low glucose (5 minutes), high glucose (10 minutes) and Exendin-4 (10 minutes)**. GFP fluorescent signals show Ca^2+^ influx into individual cells in GCaMP6f-SC-islets. Images were taken every 5 seconds for 25 minutes using 40x water objective. Video corresponds to Fig. 7C, relating to Fig. S7B.

**Video S8. Time-lapse video of live-imaged GCaMP6f-SC-islet treated with 5 ng/ml BMP4 for 11 days and exposed to low glucose (5 minutes), high glucose (10 minutes) and Exendin-4 (10 minutes)**. GFP fluorescent signals show Ca^2+^ influx into individual cells in GCaMP6f-SC-islets. Images were taken every 5 seconds for 25 minutes using 40x water objective. Video corresponds to Fig. 7D, relating to Fig. S7B.

**Table S1. Medium composition and growth factors used for co-culturing SC-islets with endothelial cells and fibroblasts in 3D fibrin gels, related to Fig. 1, S1, and 3**.

**Table S2. Medium composition and growth factors used for co-culturing SC-islets with endothelial cells and fibroblasts in microfluidic device, related to Fig. 4 and S4**.

**Table S3. Predicted cell-cell communication networks between SC-β-cells, endothelial cells, and fibroblasts, related to Fig. 7 and S7**.

(**A**) Summary of data sets used in this study.

(**B**) Predicted ligand-receptor pairs between SC-β-cells, endothelial cells, and fibroblasts.

**Table S4: Medium composition and factors used for SC-islet differentiation**.

